# Aldosterone mediates potassium-induced kidney fibrosis through tubular and mesenchymal mineralocorticoid receptor activation

**DOI:** 10.1101/2022.06.15.496280

**Authors:** Thibaut Charlemagne, Ali Sassi, Monika Gjorgjieva, Raphael Yvon, Malaury Tournier, Marylise Fernandez, Alexandra Chassot, Isabelle Roth, Christelle Veyrat-Durebex, Suresh Ramakrishnan, Joseph M. Rutkowski, Eric Feraille, Valérie Olivier

**Affiliations:** Department of Cell Physiology and Metabolism, Faculty of Medicine, University of Geneva, Geneva, Switzerland; Department of Internal Medicine, University of Texas Southwestern Medical Center, Dallas, TX, USA

## Abstract

**Background:** Mineralocorticoid receptor (MR) activation drives chronic kidney disease (CKD) progression, as evidenced by the renoprotective effects of MR antagonists in clinical trials and conditional knockout models in myeloid and vascular smooth muscle cells. Dietary potassium stimulates aldosterone secretion, yet its impact on renal fibrosis, a hallmark of CKD progression, has not been systematically investigated. We hypothesised that high dietary potassium promotes renal fibrosis through aldosterone-driven MR activation, and sought to identify the renal cell populations mediating this effect.

**Methods:** Renal fibrosis was evaluated in two murine models: unilateral ureteral obstruction (UUO, days 7–10) and inducible podocyte apoptosis (day 20), across dietary potassium concentrations ranging from 0.01% to 2%. GFR was measured by transcutaneous FITC-sinistrin clearance. Fibrosis was assessed by histology, immunofluorescence, and biochemistry. The MR dependence of the potassium effect was tested pharmacologically using finerenone and genetically using tissue-specific conditional knockout of MR in renal tubular epithelial and mesenchymal cells.

**Results:** High dietary potassium increased renal fibrosis in both models and reduced GFR by a further 50% in the podocyte apoptosis model. The pro-fibrotic effect persisted across sodium and anion variants, supporting specificity to potassium per se. Fibrosis burden correlated logarithmically with plasma aldosterone, and the pro-fibrotic effect of high potassium was attenuated by finerenone. Conditional MR deletion in tubular epithelial and mesenchymal cells independently recapitulated this protection, establishing cell-type specificity of the pathway.

**Conclusion:** High dietary potassium worsens renal fibrosis in murine models through aldosterone-dependent MR activation. This study provides the first in vivo genetic evidence that MR activation in renal tubular epithelial and mesenchymal cells can drive fibrosis, identifying a potentially actionable pathway in CKD management.

**Keypoints:** **-** High dietary potassium worsened renal fibrosis through aldosterone and mineralocorticoid receptor activation in two independent models..

**-** Tubular and mesenchymal mineralocorticoid receptor deletion independently reduced fibrosis, providing first in vivo evidence of their role

**-** Aldosterone levels correlated with fibrosis and may help identify patients most likely to respond to mineralocorticoid antagonist therapy.

## INTRODUCTION

Patients with chronic kidney disease (CKD) have historically been advised to avoid potassium-rich foods, including fruits, vegetables, nuts, and legumes, mainly for fear of hyperkalaemia. Hyperkalaemia can indeed be acutely harmful and lead to death, and is also associated with poorer chronic cardiovascular and kidney outcomes, even at mild levels in CKD patients.^1^

On the other hand, dietary approaches that reduce sodium and increase potassium intake have proven successful in managing high blood pressure and have been associated with reduced risk of stroke.^2,3^, Large-scale interventional data showing benefit in other aspects of cardiovascular risk remain limited. In light of this current evidence, the latest 2024 KDIGO recommendations on the management of CKD now call for individualised dietary potassium targets.

Aldosterone is a key mineralocorticoid hormone at the intersection of CKD and potassium homeostasis. Renin release leads to angiotensin II generation, which in turn stimulates aldosterone secretion from the adrenal cortex. Potassium directly stimulates aldosterone secretion from the adrenal cortex, independently of the renin-angiotensin axis. The hormone in turn acts on the aldosterone sensitive distal nephron (ASDN), increasing ENaC expression and activity, thereby enforcing the electrochemical gradient driving distal potassium secretion. In physiological conditions, increases in dietary potassium translate to increased kaliuresis while plasma levels remain tightly controlled. During progressive CKD, in the face of nephron loss, morphological and functional adaptations of the remaining nephrons allow for the maintenance of steady plasma ion levels until the advanced stages of the disease. When these compensatory mechanisms are overwhelmed, hyperkalaemia develops.

Beyond its canonical role in distal nephron mediated potassium regulation, mineralocorticoid receptor (MR, encoded by *Nr3c2*) activation across multiple renal compartments contributes to the development and progression of CKD. Landmark clinical trials of non-steroidal MR antagonists, particularly finerenone, have demonstrated slowing of CKD progression, providing the most direct clinical evidence for this pathway. Earlier steroidal MR antagonists such as spironolactone had shown anti-fibrotic and anti-inflammatory properties in pre-clinical studies,^1,11^ but failed to demonstrate a favourable risk-benefit ratio in CKD. In vitro and in vivo studies have further implicated aldosterone and MR activation in CKD-driving processes across multiple cell types, including podocytes, myeloid, endothelial, and vascular smooth muscle cells.

Outside the ASDN, where 11β-HSD2 is absent, MR can be activated by cortisol, which circulates at 100–1000-fold higher concentrations than aldosterone,^5^ with ligand-specific receptor conformations potentially shaping downstream signalling.^12^ Plasma aldosterone levels increase with declining kidney function,^13,14^ and may be inappropriately elevated in CKD.^15.16^ Higher levels are associated with GFR decline and progression to end-stage kidney disease.^13,14^ This findings support a prominent role of the aldosterone-MR axis in CKD pathogenesis. We therefore hypothesized that high dietary potassium could drive fibrosis, a hallmark of CKD, through aldosterone-driven MR activation

## METHODS

### Animal experiments

All experiments were performed in male mice; female reproductive anatomy complicates the UUO surgical procedure,^17^ and male sex is associated with greater susceptibility to renal fibrosis in experimental kidney injury models.^18^ All experiments were approved by the Institutional Ethical Committee of Animal Care of the University of Geneva and Cantonal veterinary authorities of the Canton of Geneva, Switzerland, and carried out in accordance with European Union Directive 2010/63/EU. Results were reported in accordance with ARRIVE guidelines.^19^ Dietary boluses were prepared from a custom purified powdered diet base (Granovit AG, Kaiseraugst, Switzerland; ref. 203600MAA05) to which sodium and potassium salts were added at the intended concentrations (% w/w). Standard diets contained 0.32% Na as sodium chloride (NS), with low (0.01%) (0.01K), normal (0.8%)(0.8K), or high (2%)(2K) potassium as potassium chloride; a medium-high potassium variant (1.3%)(1.3K) was used in models where impaired renal potassium excretion precluded the use of the 2K diet (tubular knockout and POD-ATTAC model, described below). Sodium and anion variants — including high-sodium (HS, 1.26%), low-sodium (LS, 0.06%), and potassium citrate-based diets — were also tested; full details of all diet compositions and naming conventions are provided in Supplementary Table 1. Diet (and treatment when applicable) was assigned sequentially by cage and provided ad libitum throughout.

### Nephropathy models

#### Unilateral Ureteral Obstruction Model

We performed unilateral ureteral obstruction (UUO) in 8–12-week-old C57BL/6J mice under isoflurane anaesthesia, ligating the left ureter with a 4-0 silk suture through an incision in the left flank. Diets were introduced 7 days before surgery and maintained until sacrifice, which was performed on day 7 or 10 post-UUO depending on the cohort. In a separate cohort, finerenone (Kerendia®, Bayer AG, Leverkusen, Germany), ground from tablets, was incorporated into dietary boluses at 300 ppm, approximating a dose of 10 mg/kg/day based on average daily food intake,^20,21^ from the day following UUO; drug-free boluses served as controls. Medicated and control boluses were prepared fresh daily.

#### POD-ATTAC model

POD-ATTAC transgenic mice, expressing an inducible Caspase-8 fusion protein specifically in podocytes, in a FVB background, were obtained from Dr P. Scherer (UT Southwestern Medical Center, Dallas, TX) and used as previously described.^22^ We induced podocyte apoptosis by five daily intraperitoneal injections of AP20187 (0.2 μg/g body weight; Takara Bio Inc., Kusatsu, Japan) in 8–10-week-old mice. Proteinuria was confirmed by dipstick on day 5 post-induction. Diets were started on the day of the first injection, and animals were sacrificed 20 days after the first injection.

### Genetic Knock Out models

Three tissue-specific MR (*Nr3c2*) knockout (KO) models were used, tubular/epithelial, mesenchymal and myeloid, all on a C57BL/6 background and compared against MR^lox/lox^ Cre/Cre (WT) littermates of each specific cohort.

**MR-KO_tub_** mice (MR^lox/lox^ Pax8-LC1-Cre^+^/Cre) were obtained as previously described.^23^ From 4 weeks of age, MR-KO_tub_ and WT mice received doxycycline (2 mg/mL, Thermo Scientific, USA) and sucrose (2%) in drinking water for two weeks to induce tubular *Nr3c2* deletion, followed by one week of saline supplementation to compensate for urinary sodium wasting secondary to pseudohypoaldosteronism. For the same reason, all experiments in this model were conducted on a high-sodium diet. Dietary and surgical protocols followed immediately after the induction period.

**MR-KO*_msc_*** mice (MR^lox/lox^ PDGFRβ-CreERT2/Cre) were generated by crossing MR^lox/lox^ and PDGFRβ-CreERT2/Cre mice (The Jackson Laboratory, Bar Harbor, ME, USA). MR-KO*_msc_* and WT littermates received five daily intraperitoneal injections of tamoxifen (2 mg in 200 µL castor oil, MedChemExpress, Monmouth Junction, USA) to induce mesenchymal *Nr3c2* deletion. Dietary and surgical protocols began three weeks after the first injection, as described above.

**MR-KO_mye_** mice (MR^lox/lox^ Lyz2^tm1(cre)Ifo^) with myeloid *Nr3c2* deletion, were generated by crossing MR^lox/lox^ and Lyz2^tm1(cre)Ifo^ mice (Jackson Laboratories) in which Cre is constitutively expressed under the Lyz2 promoter. Dietary and surgical protocols were applied without modification as described above.

### Sacrifice

At sacrifice, mice were anaesthetised with ketamine (100 mg/kg) and xylazine (5 mg/kg; Bayer Healthcare, Berlin, Germany), sacrificed by exsanguination, and kidneys were immediately harvested following renal pedicle clamping and placed on ice. Each kidney was bisected coronally; one half was fixed by immersion in 4% paraformaldehyde (PFA) for histological assessment, and the other was snap-frozen in liquid nitrogen for RNA and protein extraction.

### Analysis and Statistics

Histological quantification was performed using automated pixel and object classifiers in QuPath, which are inherently blinded to group allocation; final review of classifier outputs and Western blot densitometry quantification were performed without formal blinding.

Sample sizes were calculated a priori to detect a minimum between-group difference of 20%, assuming a two-sided α of 0.05 and a power of 90% (β = 0.10), based on effect sizes and variance observed in preliminary experiments. Graphs present individual data points with error bars representing means ± standard deviation. GraphPad Prism version 11.0.2 (GraphPad Software, San Diego, CA, USA) was used for statistical analyses. Welch’s t-test or one-way ANOVA were used to compare results from diet experiments according to the number of groups, with Dunnett’s post-hoc test used for comparisons against the 0.8K reference group in the latter case. When diet was tested in conjunction with a second factor (treatment or knockout), results were analysed by full-model two-way ANOVA. Post-hoc comparisons between rows and between columns only were computed using Fisher’s LSD test, treating the data as four families (one per row, one per column). R version 4.4.3 (R Core Team, 2025) was used to process blood pressure telemetry data and to perform linear regression of plasma aldosterone and cortical fibrosis area.

## RESULTS

### High potassium diet increases renal interstitial fibrosis in the unilateral ureteral obstruction model

We first investigated the effect of various levels of dietary potassium in the UUO model. Mice were fed diets containing depleted (0.01K), normal (0.8K), or high (2K) amounts of potassium. UUO was performed after seven days on diet, and mice were sacrificed seven days after surgery (Figure 1a). In a subset of mice housed in metabolic cages, 24h urinary potassium excretion increased stepwise with dietary potassium content (Figure 1b). Pooling data across experiments, plasma potassium was correlated with dietary level: the depleted group had significantly lower plasma potassium than the normal group, with a further significant though smaller difference between the normal and high potassium groups (Figure 1c).

**Figure 1.**
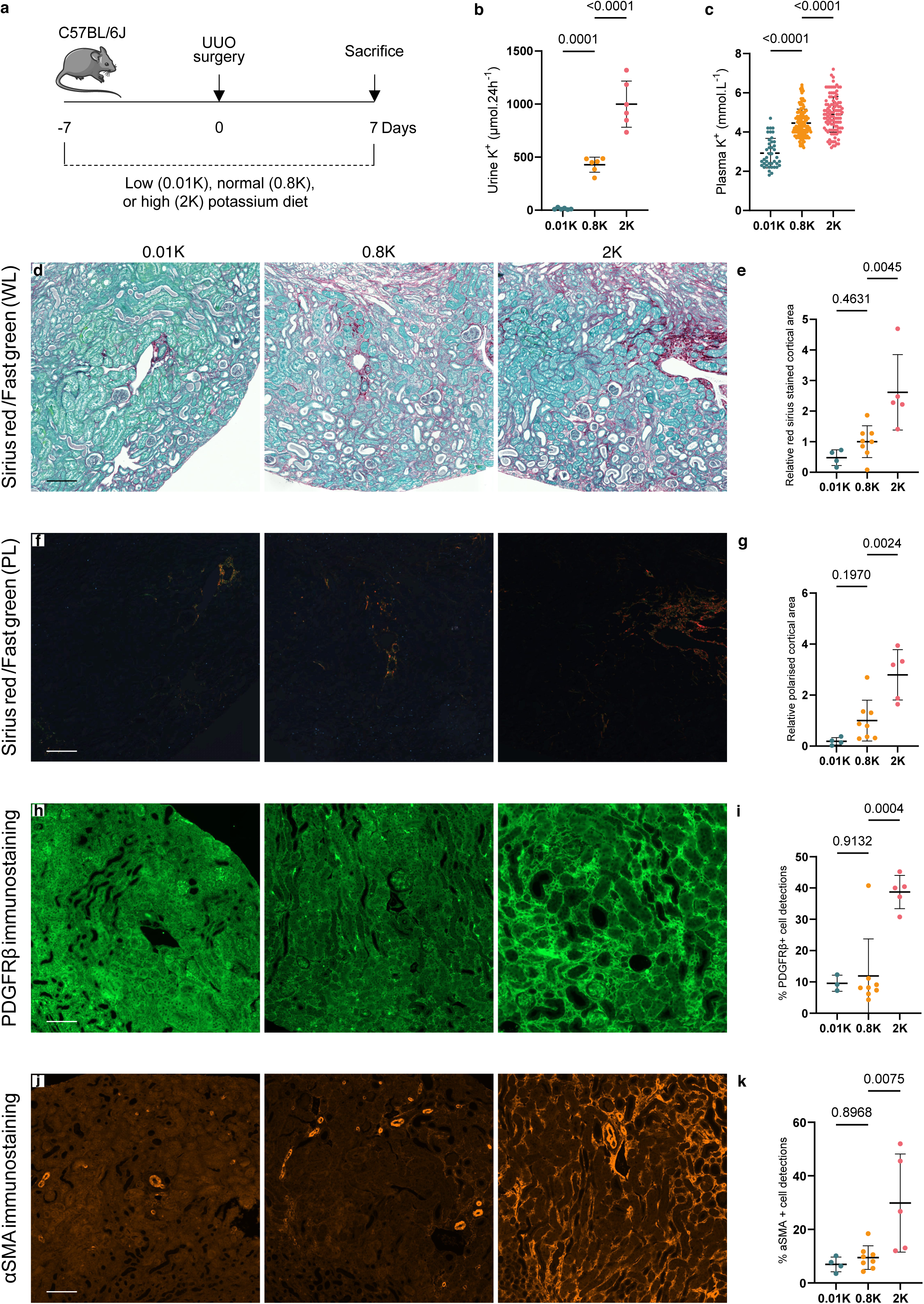
Evaluation of renal fibrosis after unilateral ureteral obstruction (UUO). **a** C57BL/6J mice were fed low (0.01K), normal (0.8K), or high (2 K) potassium diets for one week prior to UUO, then sacrificed one week post-surgery for biochemical and histological analysis. **b** In a subset of mice maintained on the same diets, urine collected via metabolic cages was analyzed by flame spectrometry (n= 0.01K:6; 0.8K:6; 2K:6; one-way ANOVA, p<0.0001). **c** Plasma potassium at sacrifice (potentiometry), pooled from several experiments (n= 0.01K: 41; 0.8K:150; 2K:111; one-way ANOVA, p<0.0001). **d, f, h, j** Representative coronal paraffin sections (left to right: 0.01K, 0.8K, 2K) stained with Sirius red and imaged under white light (WL, **d**) and polarized light (PL, **f**), or immunostained for PDGFRβ (**h**) and αSMA (j); scale bar 100 µm (n=0.01K: 4; 0.8K: 8; 2K: 5; one animal excluded from panels **h–i** in which reliable classifier performance could not be achieved were excluded from quantification.). Whole slides were digitized and quantified using a high-resolution scanner. **e** Quantification of cortical Sirius red area (WL), normalized to the 0.8K mean (one-way ANOVA, p=0.0018). **g** Quantification of cortical Sirius red polarized area (PL), normalized to the 0.8K mean (one-way ANOVA, p=0.0005). i Quantification of PDGFRβ-positive cells (one-way ANOVA, p=0.0004). **k** Quantification of αSMA-positive cells (one-way ANOVA, p=0.0061).

We then assessed interstitial fibrosis across these diets. Sirius red staining on paraffin sections from obstructed kidneys, observed under white (Figure 1d) and polarised light (Figure 1f), showed that fibrosis was associated with dietary potassium content and was higher in the 2K than the 0.8K group (Figures 1e, 1g). Fibronectin protein abundance was also associated with dietary potassium, with a more pronounced increase from 0.8K to 2K (Supplementary Figure 1a). PDGFRβ- and αSMA-positive cell counts in the cortical interstitium increased with dietary potassium, especially from 0.8K to 2K, and were particularly abundant between tubules in the 2K group (Figures 1,h-k), consistent with mesenchymal cell and myofibroblast expansion.

These data suggest that higher potassium intake leads to increased renal fibrosis in a UUO nephropathy model.

### High potassium diet increases fibrosis in a glomerular model of kidney injury

To strengthen these findings in a second model and explore potential functional consequences, we studied the effect of dietary potassium in the POD-ATTAC mouse model, in which podocyte apoptosis can be induced by injection of AP20187 (Supplementary Figure 2a). 0.01K, 0.8K, and 1.3K diets were introduced on the day of the first injection and maintained until sacrifice 20 days later (Figure 2a). In a subset of mice housed in metabolic cages, urinary potassium excretion was correlated with diet (Figure 2b). Pooling data across experiments, plasma potassium was correlated with dietary potassium intake, confirming effective dietary potassium separation and altered potassium handling in this model (Figure 2c).

**Figure 2.**
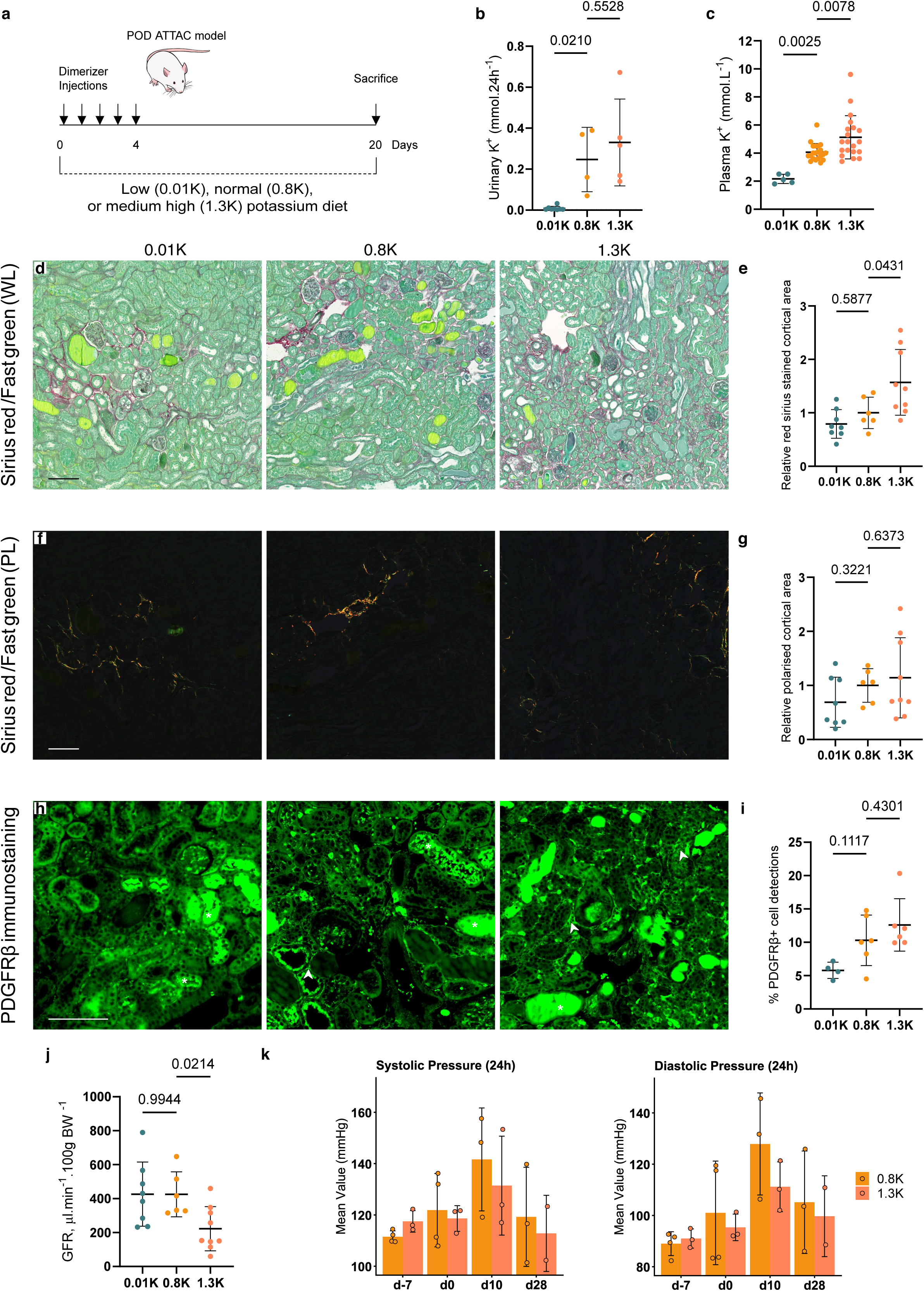
Evaluation of renal fibrosis after podocyte apoptosis induction (POD-ATTAC). **a** POD-ATTAC mice were placed on low (0.01 K), normal (0.8 K), or medium-high (1.3 K) potassium diets on the day of the first of five daily injections to induce podocyte apoptosis, and sacrificed 20 days later for biochemical and histological analysis. **b** Urinary potassium collected via metabolic cages from a subset of mice under the same protocol, measured by flame spectrometry (n= 0.01K:8; 0.8K:4; 1.3 K:5; one-way ANOVA, p=0.0021). **c** Plasma potassium at sacrifice (potentiometry), pooled from several experiments (n= 0.01K:5; 0.8K:21; 1.3K:20; one-way ANOVA, p<0.0001). **d, f, h** Representative coronal paraffin sections (left to right: 0.01K, 0.8K, 1.3K) stained with Sirius red under white light (WL, **d**) and polarized light (PL, **f**), or immunostained for PDGFRβ (**h**); scale bar 100 µm. In panel **h**, arrowheads indicate PDGFRβ-positive interstitial cells of interest; asterisks indicate non-specific fluorescence from tubular casts excluded from quantification. Whole slides were digitized and quantified using a high-resolution scanner. (d,f n= 0.01K:8; 0.8K:6; 1.3K:9) **e** Quantification of cortical Sirius red area (WL), normalized to the 0.8K mean (one-way ANOVA, p=0.0051). **g** Quantification of cortical Sirius red polarized area (PL), normalized to the 0.8K mean (one-way ANOVA, p=0.2717). **i** Quantification of PDGFRβ-positive interstitial cells (Samples in which reliable classifier performance could not be achieved were excluded from quantification. n=0.01K: 4; 0.8K:6; 1.3K: 6; one-way ANOVA, p=0.0281). **j** GFR measured by transdermal FITC-sinistrin clearance, expressed in µL per 100 g body weight (same n as in panels d-g, one-way ANOVA, p=0.0084). **k** Blood pressure was recorded over 24 hours in a subset of mice implanted with PA-C10 telemetry devices at four time points: before dietary intervention (day −7), after one week on diet and immediately prior to the first injection (day 0), during the acute injury phase (day 10), and during the chronic phase (day 28). Mean 24-hour systolic and diastolic blood pressure were recorded in the 0.8K and 1.3K groups (n= 0.8K:4;1.3K:3 at baseline; one animals per group died during the experimental period).

Kidney interstitial fibrosis was assessed with the same methodology as in the UUO model. Sirius red staining increased, with the increase in dietary potassium, under white (Figures 2d, 2e) but not polarised light (Figures 2f, 2g). This may stem from a lower level of fibrosis overall in this model. Fibronectin protein abundance increased with dietary potassium (Supplementary Figure 2b), as did PDGFRβ-positive interstitial cell counts (Figures 2h, 2i); Therefore, similarly to the UUO model, ECM-producing cells and ECM accumulation increased with dietary potassium.

Elevated dietary potassium was also associated with lower GFR on day 19, with mean body weight-normalised GFR approximately 50% lower in the 1.3K group (0.8K: 425.6 ± 132.7; 1.3K: 217.3 ± 124.1 µL·min¹·100g BW¹) (Figure 2j). This was not explained by haemodynamic changes. Blood pressure was comparable in the high potassium group and rose less sharply in response to podocyte injury (Figure 2k, Supplementary figure 2c).

### High potassium-induced interstitial fibrosis persists under high sodium condition

In the context of high blood pressure management, the effect of dietary potassium is substantially influenced by sodium content, raising the question of whether the observed pro-fibrotic effect of potassium depended on its co-administration with sodium chloride. To address this, we tested 0.01K, 0.8K, and 2K diets on a high-sodium background (HS, 1.26% Na) in the UUO model, following the same protocol as described above.

Under HS conditions, the increase in dietary potassium was still associated with a progressive increase in cortical Sirius red staining under both white light (Figure 3a, b) and polarised light (Figure 3c, d). Fibronectin protein abundance also increased with dietary potassium (Figure 3e), most markedly in the 2K group. The pro-fibrotic effect of potassium thus persisted in a high-sodium environment, arguing that the pro-fibrotic effect of high potassium is independent of sodium content.

**Figure 3.**
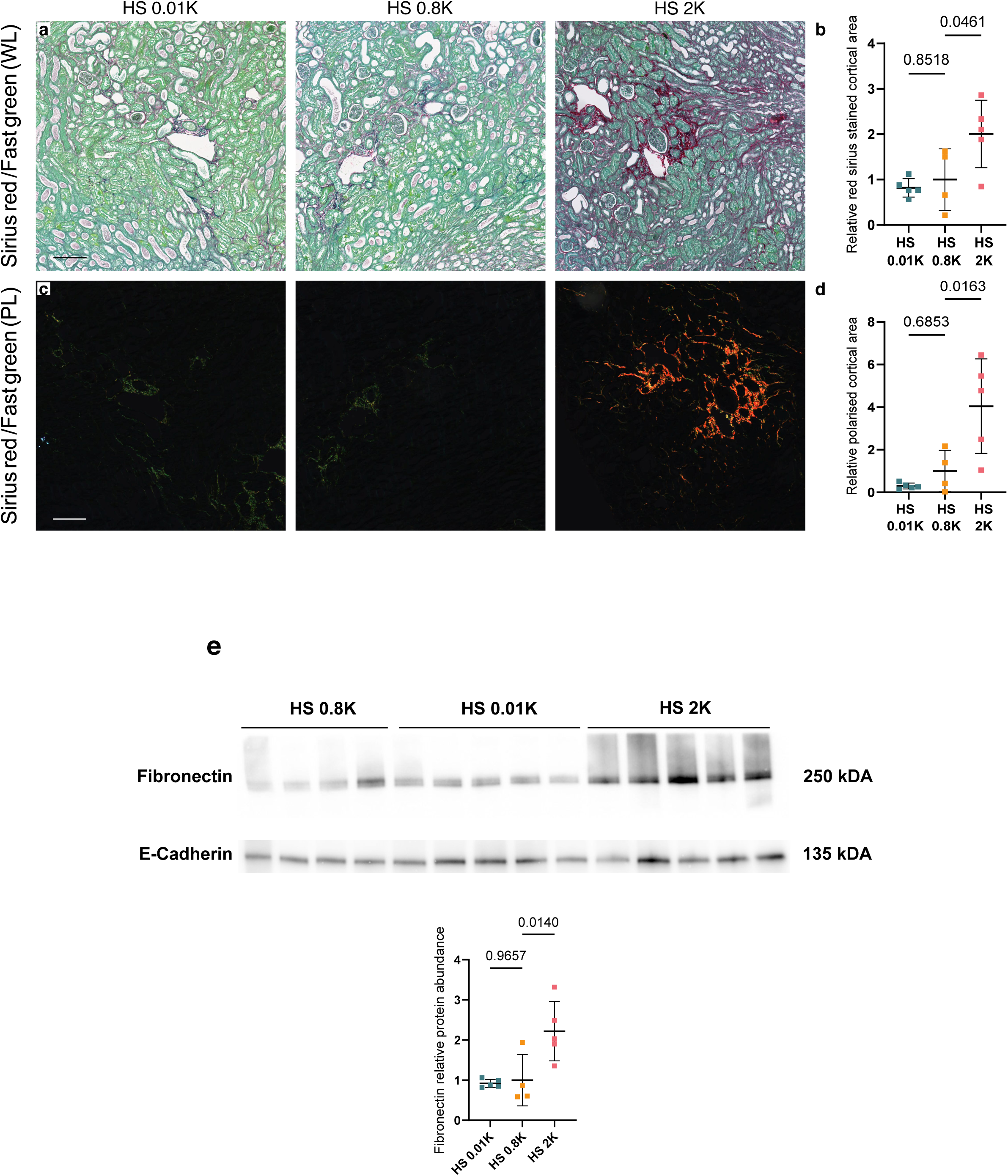
Effect of high sodium on the potassium–fibrosis relationship. Mice were fed 0.01K, 0.8K, or 2K diets in conjunction with high sodium (HS, 1.26 Na) following the same UUO time course as in Figure 1 (for all panels, n= 0.01K: 5; 0.8K: 4; 2K: 5). Representative coronal paraffin sections stained with Sirius red and imaged under white light (WL; **a**) and polarized light (PL; **c**); scale bar 100 µm. **b** Quantification of cortical Sirius red area (WL), normalized to the HS-0.8K mean (one-way ANOVA, p=0.0182).**d** Quantification of cortical Sirius red polarized area (PL), normalized to the HS-0.8K mean (one-way ANOVA, p=0.0040). **e** *Top*: Representative Western blots for fibronectin and E-cadherin. *Bottom*: Fibronectin band intensity normalized to E-cadherin and expressed relative to the HS-0.8K group mean (one-way ANOVA, p=0.0065).

Substituting chloride with citrate as the potassium counter-ion similarly failed to abolish fibrosis in either the UUO or POD-ATTAC model (Supplementary Figure 3), further supporting specificity to potassium per se.

### High potassium-induced interstitial fibrosis is dependent on mineralocorticoid receptor activation

The pro-fibrotic effect of a high potassium diet points to the potential involvement of aldosterone. We measured aldosterone across a range of sodium, potassium, and model conditions (Figure 4a). Aldosterone concentration rose progressively across potassium levels in UUO high-salt and normal-salt conditions, as well as in the POD-ATTAC model. We identified a linear correlation between the logarithm of plasma aldosterone and cortical Sirius red area under white light, which, while not explaining all the variability, was highly significant (p=0.0005, r²=0.2062) (Figure 4b).

**Figure 4.**
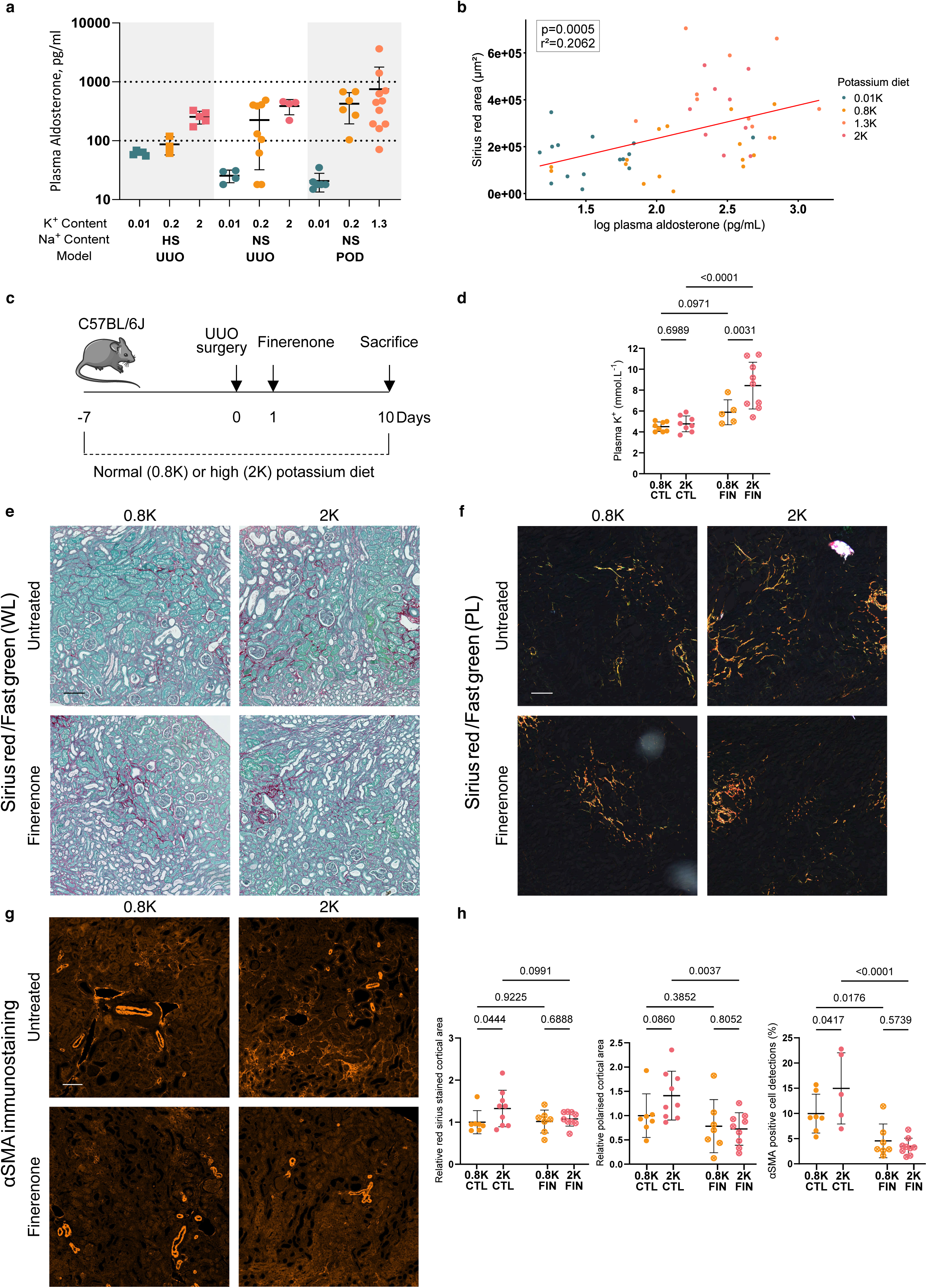
Association of dietary potassium, aldosterone, and renal fibrosis. **a** Plasma aldosterone (radioimmunoassay, pg/mL, log scale) across sodium and potassium diet conditions and models No formal statistical comparisons were performed between subgroups given the small and unequal sample sizes within each condition. (n: HS UUO 0.01K=5; 0.8K=3; 2K=5; NS UUO 0.01K=4; 0.8K=9; 2K=4; POD-ATTAC 0.01K=6; 0.8K=6; 1.3K=11) **b** Univariate linear regression of cortical Sirius surface against the log of plasma aldosterone (n=53 pooled across all cohorts and conditions shown in panel a, log-aldosterone: p=0.0005, adjusted r²=0.2062); potassium diet groups are colour-coded without adjustment. **c** C57BL/6J mice were fed normal (0.8K) or high (2K) potassium chloride diets one week prior to UUO. From the day following surgery, mice received either finerenone-supplemented (FIN) or drug-free control (untreated, CTL) dietary boluses. Mice were sacrificed 10 days post-UUO. **d** Plasma potassium at sacrifice (potentiometry; n= 0.8K CTL :7; 2K CTL :8; 0.8K FIN :5; 2K FIN:9; two-way ANOVA: p interaction=0.0399, p treatment<0.0001, p diet=0.0125, assay failure in some samples). **e, f, g** Representative coronal paraffin sections stained with Sirius red under white light (WL, **e**) and polarized light (PL, **f**) (n= 0.8K CTL :7; 2K CTL :9; 0.8K FIN :7; 2K FIN:9 for **e** and **f**) or immunostained for αSMA (**g**)( Samples in which reliable classifier performance could not be achieved were excluded from quantification. n= 0.8K CTL :7; 2K CTL :5; 0.8K FIN :7; 2K FIN:9); scale bar 100 µm. Whole slides were digitized (**h**). *Left*: Quantification of WL Sirius red area, normalized to the 0.8K control mean (two-way ANOVA: p interaction=0.2395, p treatment=0.3006, p diet=0.0869). *Middle*: Quantification of PL Sirius red area, normalized to the 0.8K control mean (two-way ANOVA: p interaction=0.1625, p treatment=0.0101, p diet=0.2884). *Right*: Quantification of αSMA-positive cells (two-way ANOVA: p interaction=0.0567, p treatment<0.0001, p diet=0.2203).

To test whether this aldosterone-dependent pathway was causally involved in the pro-fibrotic effect of high potassium, we used the selective MR antagonist (MRA) finerenone. Finerenone was incorporated into dietary boluses from the day following UUO (Figure 4c). Plasma potassium increased markedly with finerenone, especially in conjunction with the 2K diet, confirming treatment and dietary compliance (Figure 4d). Finerenone-treated mice showed reduced obstructed kidney weight and smaller coronal section area, with increased cell density suggesting cellular volume contraction rather than cell loss; this was not influenced by dietary potassium (Supplementary Figure 4). Because kidney section area was influenced by finerenone treatment, raw cortical Sirius red area was used for quantification rather than a ratio to total section area. Although white light Sirius Red staining showed no significant overall effect, post-hoc pairwise comparisons indicated a diet-associated increase in untreated animals that was not observed in the finerenone group (Figure 4e, 4h left). Polarised light signal was reduced by finerenone (Figure 4f, 4h centre). The diet effect did not reach statistical significance in this analysis, potentially reflecting the influence of a single divergent data point on the overall variance. Fibronectin protein abundance (Supplementary Figure 5), and αSMA-positive cell counts (Figures 4g, 4h right) showed the same pattern: increase with the 2K diet in untreated animals and attenuation by finerenone. As independent validation, spironolactone similarly decreased interstitial fibrosis in 1.3K-fed POD-ATTAC mice (Supplementary Figure 6).

### Mesenchymal and tubular epithelial MR KO, but not myeloid, reduce high potassium-induced kidney fibrosis

Given the MR dependence of the potassium-induced fibrotic process, we sought to identify the specific cell populations mediating this effect by systematically evaluating tissue-specific MR knockout models (KO). Myeloid MR ablation did not meaningfully affect the fibrotic response to dietary potassium (Supplementary Figure 7).

By contrast, the tubular epithelial specific MR KO proved protective in the UUO model. MR^lox/lox^ Pax8-LC1-Cre/Cre (MR-KO_tub_) and WT littermates received doxycycline for KO induction and were maintained on a high-salt (HS) diet with normal (HS 0.8K) or medium-high potassium (HS 1.3K) to manage induced pseudohypoaldosteronism before undergoing UUO (Figure 5a). Successful recombination was confirmed by reduced *Nr3c2* expression in contralateral non obstructed kidney tissue (Supplementary Figure 8a).

**Figure 5.**
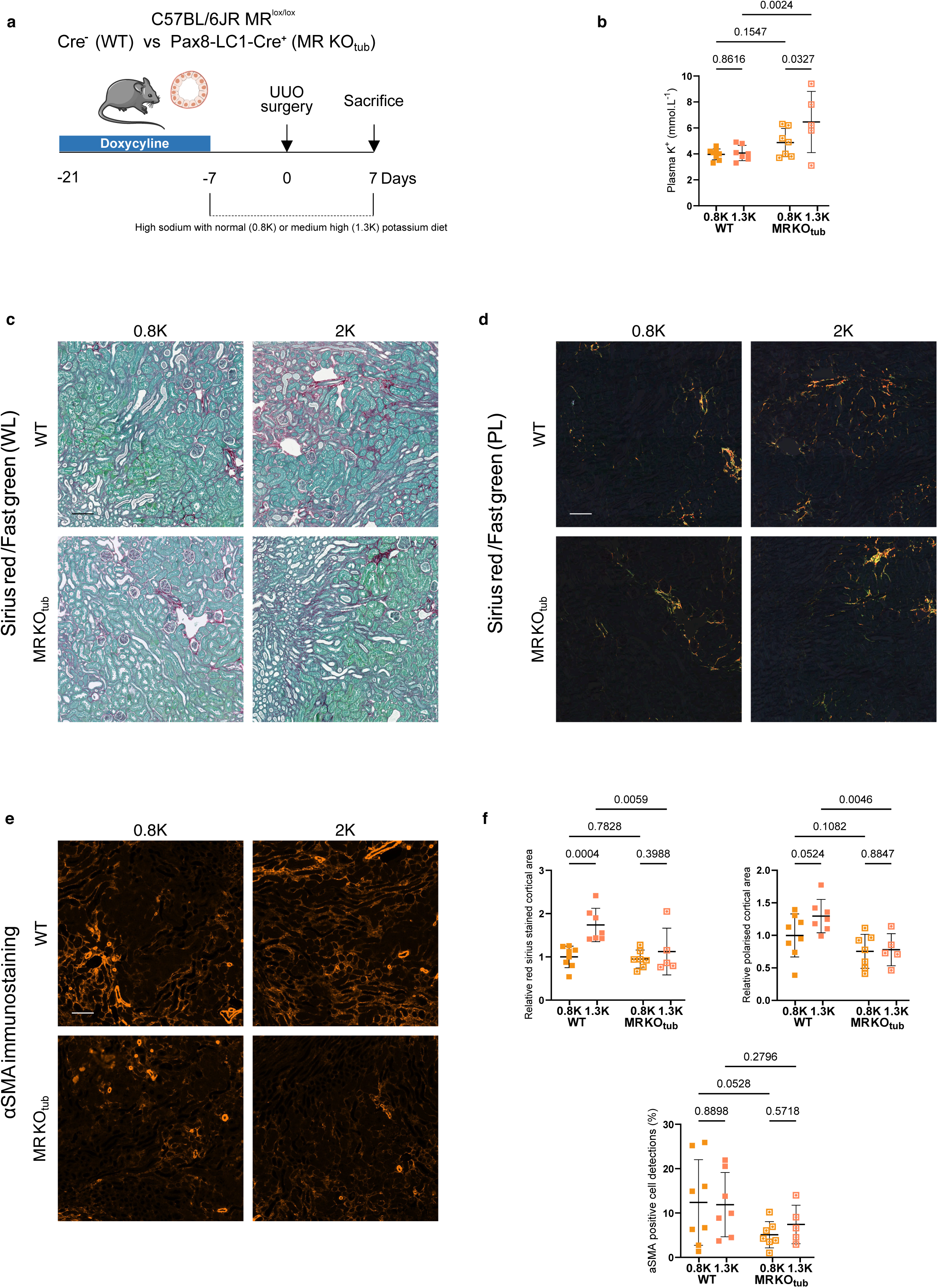
Effect of tubular mineralocorticoid receptor knockout (KO) on renal fibrosis. **a** MR^lox/lox^ Pax8-LC1-Cremice (MR-KO_tub_) and their MR^lox/lox^ Cre littermates (WT) received doxycycline for two weeks to induce tubular *Nr3c2* deletion, with concurent saline supplementation. All animals were maintained on a high-sodium diet throughout to compensate for pseudohypoaldosteronism. UUO was performed three weeks after the start of induction, with HS 0.8K or HS1.3K diets introduced one week before surgery and sacrifice 7 days post-procedure. (for all panels, n= HS 0.8K WT :8; HS 1.3K WT :7; HS 0.8K MR-KO_tub_:7; HS 1.3K MR-KO_tub_:5) **b** Plasma potassium at sacrifice (potentiometry; two-way ANOVA: p interaction=0.1263, p genotype=0.0018, p diet=0.0818). **c, d, e** Representative coronal paraffin sections stained with Sirius red under white light (WL, **c**) and polarized light (PL, **d**), or immunostained for αSMA (**e**); scale bar 100 µm. Whole slides were digitized and quantified (**f**). *Top left*: Quantification of WL Sirius red area, normalised to the 0.8K WT mean (two-way ANOVA: p interaction=0.0481, p genotype=0.0220, p diet=0.0026). *Top right*: Quantification of PL Sirius red area, normalised to the 0.8K WT mean (two-way ANOVA: p interaction=0.2258, p genotype=0.0021, p diet=0.1566). *Bottom*: Quantification of αSMA-positive cells (two-way ANOVA: p interaction=0.6063, p genotype=0.0396, p diet=0.7392).

The mesenchymal specific MR KO also proved protective in the UUO model. MR^lox/lox^ PDGFRβ-CreERT2/Cre (MR-KO*_msc_*) and WT littermates received tamoxifen for KO induction and were placed on normal salt with 0.8K or 2K diets before UUO (Figure 6a).

**Figure 6.**
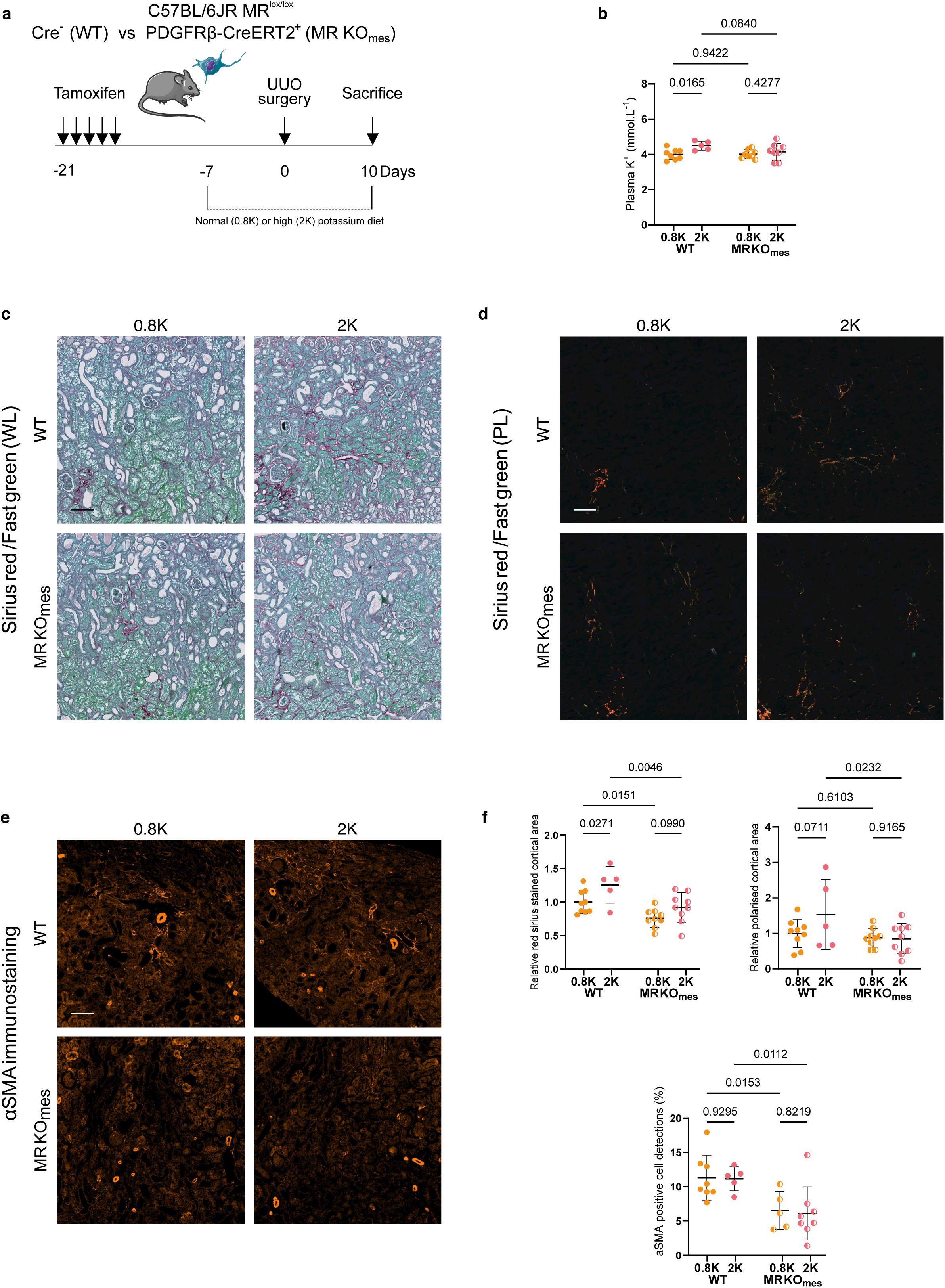
Effect of mesenchymal mineralocorticoid receptor knockout (KO) on renal fibrosis. **a** C57BL/6J MRlox/lox PDGFRβ-CreERT2 mice (MR-KO_msc_) and their MRlox/lox Cre littermates (WT) were induced with tamoxifen three weeks prior to UUO and placed on 0.8K or 2K diets one week before surgery, then sacrificed 10 days post-procedure. **b** Plasma potassium at sacrifice (potentiometry; n= 0.8K WT :8; 2K WT :5; 0.8K MR-KO_msc_:8; 2K MR-KO_msc_:8, assay failure in some samples; two-way ANOVA: p interaction=0.1734, p genotype=0.2038, p diet=0.0209). **c, d, e** Representative coronal paraffin sections stained with Sirius red under white light (WL, **c**) and polarized light (PL, **d**)( n= 0.8K WT :9; 2K WT :5; 0.8K MR-KO_msc_:9; 2K MR-KO_msc_:9), or immunostained for αSMA (**e**) (Samples in which reliable classifier performance could not be achieved were excluded from quantification n= 0.8K WT :8; 2K WT :5; 0.8K MR-KO_msc_:5; 2K MR-KO_msc_:8); scale bar 100 µm. Whole slides were digitized and quantified (**f**). *Top left*: Quantification of WL Sirius red area, normalized to the 0.8K WT mean (two-way ANOVA: p interaction=0.5023, p genotype=0.0004, p diet=0.0075). *Top right*: Quantification of PL Sirius red area, normalized to the 0.8K WT mean (two-way ANOVA: p interaction=0.1445, p genotype=0.0389, p diet=0.1833). *Bottom*: Quantification of αSMA-positive cells (two-way ANOVA: p interaction=0.9229, p genotype=0.0009, p diet=0.8245).

Recombination was confirmed by the presence of a shortened *Nr3c2* exon 3 amplicon in fibroblast-rich ear pinnae (Supplementary Figure 9a).

MR-KO_tub_ mice showed elevated plasma potassium, especially in the 1.3K subgroup, consistent with impaired kaliuresis following tubular MR KO, mirroring the effect of finerenone (Figures 5b). MR-KO*_msc_* did not affect plasma potassium (Figure 6b). Similarly to pharmacological MR inhibition, inducible ablation of MR in tubular epithelial cells reduced obstructed kidney size (Supplementary Figure 8b). In both models, Sirius red staining under white light was increased by high dietary potassium and attenuated by KO on the main effect analysis, with post-hoc testing showing loss of the diet effect in KO animals (Figures 5c, 5f top left, 6c, 6f top left). Polarised light signal was similarly reduced by both KOs (Figures 5d, 5F top right, 6d, 6f top right). αSMA-positive cell counts were attenuated in both MR-KO_tub_ and MR-KO*_msc_*mice (Figures 5e, 5f bottom, 6e, 6f bottom). Fibronectin protein abundance was also reduced by MR-KO_tub_ (Supplementary Figure 8c, d). Results were not significant in the MR-KO*_msc_* cohort (Supplementary figure 9b), possibly reflecting technical limitations; however, mRNA expression of Collagen1α1, PDGFRβ, and αSMA increased with the 2K diet and were reduced or tended to be reduced by MR-KO*_msc_*, consistent with diet-driven ECM accumulation and mesenchymal cell activation attenuated by the knockout (Supplementary Figure 9c).

Therefore, while myeloid MR did not affect the fibrotic response to dietary potassium, both tubular and mesenchymal-derived MR promote fibrosis in the UUO model.

## DISCUSSION

Our data demonstrate a pro-fibrotic effect of high dietary potassium across two mechanistically distinct models of kidney injury, extending a prior report in the rat 5/6 nephrectomy model.^24^ The persistence of this effect across high sodium content and anion substitution conditions supports specificity to potassium per se. These findings contrast with observational human data collated by Picard et al. and discussed at the 2021 KDIGO controversies conference,^25,26^ where higher dietary potassium intake was associated with slower CKD progression. Two factors help reconcile this discrepancy.

First, the potassium dose is critical. Standard laboratory chow approximates the WHO-recommended intake of 3.5 g/day when normalised to caloric intake,^27^ whereas intakes in CKD cohorts are substantially lower, with a median of 1,8 g/day reported by Suenaga et al.^28^ The diets used here therefore represent the extreme upper range of human exposure, more comparable to potassium supplementation than to typical dietary variation.

Second, bioavailability substantially modulates the relationship between dietary and plasma potassium.^29^ The high bioavailability of our dietary boluses likely explains the marked elevation of plasma potassium, which more closely resembles levels during a potassium supplementation trial in CKD, ^30^ than those across dietary quartiles in population studies.^25,28^ Although hyperkalaemia may contribute to adverse outcomes in CKD,^1^ fibrosis was attenuated in high-potassium MRA and MR-KO_tub_ groups, despite extreme plasma potassium levels, suggesting that hyperkalaemia in this context may reflect the associated aldosterone response rather than independently driving fibrosis.

Aldosterone levels correlated logarithmically with interstitial fibrosis across models, consistent with prior experimental and clinical observations.^13^D^24^ Potassium supplementation increases plasma aldosterone without increasing renin activity,^30^ consistent with direct adrenal stimulation. Subclinical primary aldosteronism, distinct from overt hyperaldosteronism, is increasingly recognised as an independent risk factor for CKD progression,^31^ raising the possibility that sustained potassium-driven aldosterone elevation could perpetuate renal fibrosis without frank hyperkalaemia. Supporting this, CRIC cohort data showed that higher aldosterone was associated with greater urinary potassium excretion but not with the highest plasma potassium values,^13^ suggesting that high dietary potassium may be managed through enhanced kaliuresis at the cost of sustained aldosterone elevation and its downstream pro-fibrotic consequences.

Independently of potassium, recent landmark trials of finerenone in diabetic^8^ and non-diabetic^9^ CKD have firmly established the therapeutic relevance of MR antagonism. Consistent with this, finerenone reduced fibrosis and suppressed the pro-fibrotic effect of high dietary potassium, supporting MR dependence. Finerenone-treated animals showed a reduction in obstructed kidney weight, previously reported across several models,^20,32,33^ and a qualitatively similar reduction was observed in MR-KO_tub_ mice, possibly reflecting a comparatively high effective drug concentration in our model.

MR is expressed across multiple renal cell types, but their relative contributions to fibrosis remain incompletely defined. Myeloid MR deletion proved protective in models of anti-glomerular basement membrane glomerulonephritis^34^ and ischaemia-reperfusion injury,^35^ but not in mice with concomitant hypertension and diabetes,^36^ compatible with our own negative findings. Smooth muscle cell and parietal epithelial cell MR deletion have shown benefit in cyclosporine nephropathy^37^ and crescentic glomerulonephritis^38^ respectively, illustrating the context-dependence of cell-type specific MR effects.

Injured proximal tubular cells and activated myofibroblasts have recently been proposed as converging drivers of renal fibrosis and functional decline,^39^ and single-cell multi-omics data in a rat DOCA/salt model identified phenotype-driving alterations in proximal tubular gene expression under finerenone treatment.^40^ Here, we provide the first in vivo evidence that MR deletion in tubular epithelial cells attenuates renal fibrosis with reduced myofibroblast activation and ECM deposition. Earlier in vitro studies described aldosterone-driven epithelial-mesenchymal transition, a framework now largely abandoned in vivo. It is more likely that injured tubular cells promote fibrosis through paracrine effects on myofibroblasts and inflammatory cells.^39D41^ and as suggest by our data, activation of MR could shape this signalisation.

Our results also establish a role for mesenchymal MR in renal fibrosis, with attenuated ECM accumulation and myofibroblast activation in the PDGFRβ-specific MR-KO. Prior evidence was limited to in vitro studies showing that aldosterone promotes fibroblast proliferation and ECM production.^42–44^ Myofibroblasts are widely considered to arise from resident pericytes and fibroblasts,^39,41^ both falling within the PDGFRβ-expressing lineage targeted by our KO. These results demonstrate that MR activation governs the pro-fibrotic fate of mesenchymal cells in this context, warranting further characterisation across these lineages.

Importantly, fibrosis attenuation in both KO models occurred in the context of potassium-driven aldosterone excess, directly linking dietary potassium, aldosterone secretion, and MR activation across two distinct renal cell lineages. Whether aldosterone activates independent pro-fibrotic pathways in parallel across these compartments or acts at successive levels of a self-reinforcing loop remains to be determined.

The translatability of these findings warrants caution. Our high-bioavailability potassium boluses differ substantially from plant-based dietary potassium and from the intake range typical of CKD populations. Nonetheless, recent evidence suggests a U-shaped relationship between potassium intake and both CKD progression^28^ and blood pressure,^45^ with outcomes modelled to worsen beyond approximately 2.3 and 3.5 g/day respectively. A negative association between dietary potassium and measured GFR was similarly observed in the companion clinical study to this work and in a healthy high-intake population cohort.^46^ These apparent discrepancies with larger observational studies,^25,26^ likely reflect differences in baseline intake, potassium bioavailability, and aldosterone status, as well as CKD stage (a heterogeneity acknowledged by Picard et al. themselves, who noted that adverse or null associations were more prominent at later stages of disease²).

These findings are not incompatible with the benefits of plant-based diets in CKD, whose potassium carries substantially lower bioavailability,^29^ and whose renoprotective effects are mediated through multiple mechanisms beyond potassium content. They do support the growing case for aldosterone as a driver of CKD progression.^13D31^ As the range of approved therapies in CKD continues to expand, encompassing SGLT2 inhibitors, GLP-1 agonists, MR antagonists, and emerging aldosterone synthase inhibitors, the question of how to individualise treatment selection arises. Our data raise the possibility that aldosterone monitoring could serve as a stratification tool to identify patients most likely to benefit from MR antagonism or aldosterone synthase inhibition, particularly those in whom dietary potassium or other stimuli sustain elevated aldosterone levels.

The present study employs two acute injury models — obstructive nephropathy, propagating from the medulla outward, and podocyte apoptosis, initiating glomerular injury from the filtration unit — which, while mechanistically distinct and mutually reinforcing, do not establish whether the same pathway operates in slowly progressive CKD. Their short experimental timeframe was however instrumental in enabling systematic screening of multiple tissue-specific KO models, a level of mechanistic resolution difficult to achieve in longer-running models, and future studies can now be rationally focused on the tubular and mesenchymal compartments identified here. The downstream molecular mechanisms by which MR activation drives fibrosis in these cells remain to be defined. Finally, the high bioavailability of our dietary boluses limits direct comparability with plant-based dietary approaches, though this same feature allowed potassium to be studied in isolation.

In conclusion, we demonstrate that high intake of highly bioavailable potassium increases renal fibrosis across two distinct models of kidney injury, an effect correlated with aldosterone elevation and dependent on MR activation, as evidenced by its attenuation with finerenone. We further provide the first in vivo genetic evidence that MR deletion in tubular epithelial and mesenchymal cells independently attenuates fibrosis, establishing these compartments as effectors of the aldosterone-MR fibrotic axis. These findings raise the possibility that aldosterone monitoring could serve as a stratification biomarker to guide dietary and pharmacological interventions targeting MR activation in CKD.

## Disclosure

All the authors declared no competing interests.

## Author Contributions

**Conceptualization:** TC, EF,VO

**Formal analysis:** TC, AS, MG, EF, VO

**Funding acquisition:** EF

**Investigation:** TC, AS, RY, MT, MF, AC, IR, CV-D, SR, VO

**Methodology:** TC, MT, MF, IR, JMR, VO

**Supervision:** EF, VO

**Visualization:** TC, VO

**Writing – original draft:** TC, EF, VO

**Writing – review & editing:** TC, AS, MG, CV-D, SR, JMR, EF, VO

## Aknowledgements

We thank Prof. G. Schütz for providing the MR^lox/lox^ mouse line, Prof. E. Hummler for the Pax8-LC1-Cre mouse line, and Dr P. Scherer (UT Southwestern Medical Center, Dallas, TX) for generously sharing POD-ATTAC transgenic mice. We thank the laboratory of renal physiology, Centre de Recherche des Cordeliers, INSERM UMR_S 1138, CNRS ERL 8228, Sorbonne Université, Paris, France, for technical assistance with urinary potassium measurements by flame photometry, and Dr Maillard (Centre Hospitalier Universitaire Vaudois, Lausanne, Switzerland) for aldosterone radioimmunoassay measurements. We thank the Zootechnical, Bioimaging and Histology Core Facilities at the University of Geneva for their technical support.

## Data Availability Statements

Original data generated for the study will be made available upon reasonable request to the corresponding author. Data Type: Image Data; Raw Data/Source Data.

## Declaration of generative AI and AI-assisted technologies in the manuscript preparation process

During the preparation of this work the authors used Claude Sonnet 4.6 (Anthropic) to assist with manuscript editing, and language refinement, including rephrasing of figure legends and text sections. All content was reviewed, edited, and approved by the authors, who take full responsibility for the published article.

