## Supplementary methods and figures for "Aldosterone mediates potassium-induced kidney fibrosis through tubular and mesenchymal mineralocorticoid receptor activation"

**Charlemagne T, et al**

Table of contents:

| | |
|---|---|
| Supplementary methods | Page 2 |
| Supplementary Table 1 | Page 10 |
| Supplementary Table 2 | Page 11 |
| Supplementary Figure 1 | Page 12 |
| Supplementary Figure 2 | Page 13 |
| Supplementary Figure 3 | Page 14 |
| Supplementary Figure 4 | Page 15 |
| Supplementary Figure 5 | Page 16 |
| Supplementary Figure 6 | Page 17 |
| Supplementary Figure 7 | Page 18 |
| Supplementary Figure 8 | Page 19 |
| Supplementary Figure 9 | Page 20 |
| Supplementary Figure 10 | Page 21 |
| Supplementary Figure 11 | Page 22 |
| Supplementary Figure 12 | Page 23 |

### Supplementary methods

*Extended animal methods*

All animals were housed in the animal facilities of the University of Geneva under conventional housing conditions. Health monitoring was performed quarterly for key pathogens and annually for others, in accordance with FELASA recommendations, by the Service de Zootechnie of the University of Geneva. The facility is positive for *Helicobacter hepaticus*, *H. bilis*, and *H. typhlonicus*, pinworms (*Syphacia obvelata*, *Aspicularis tetraptera*), apathogenic protozoa (*Chilomastix* sp., *Entamoeba* sp., *Hexamastix* sp., *Trichomonas* sp.), and sporadic detection of *Rodentibacter* and Mouse Hepatitis Virus and Mouse Norovirus. All other monitored pathogens were absent. Animals were housed in open cages until early 2025 and in individually ventilated cage systems thereafter. Animal facilities maintained a temperature of 21 ± 1°C and relative humidity of 50± 20%, with a 12-hour light/dark cycle.

Non-genetically modified C57BL/6 mice were obtained from Charles River France, while mineralocorticoid receptor (MR) knockout (KO) lines were bred in-house on a C57BL/6 background. The $MR^{lox/lox}$ mouse line, in which a loxP site introduced 170 bp upstream of exon 3 of *Nr3c2* and a second loxP site flanking a neomycin resistance–thymidine kinase cassette allow Cre-mediated excision of exon 3 encoding the first zinc finger of the DNA-binding domain, has been described previously.[S1] $MR^{lox/lox}$ mice were crossed with mice expressing tissue-specific Cre recombinases to generate tissue-specific KO lines. Genotyping was performed by PCR on genomic DNA from phalanx biopsies; primers are listed in Supplementary Table 2

POD-ATTAC mice were also bred in-house. Genotyping was performed by PCR on phalanx biopsies as described above; primers are listed in Supplementary Table 2. These mice express a FKBP/Caspase-8 fusion protein under the control of the *NPHS2* (podocin) promoter. Kidney fibrosis was induced by once-daily intraperitoneal injection of AP20187 (0.2 µg/g body weight; Takara Bio Inc., Kusatsu, Japan), an FK506-derived compound, for 5 days in 8–10-week-old male transgenic mice. This compound induces dimerization of the fusion protein and Caspase-8 activation, resulting in podocyte-specific apoptosis and glomerular proteinuria.[22]

GFR was measured by transcutaneous FITC-sinistrin clearance. Mice were anaesthetised with isoflurane and a miniaturised imager device (Mannheim Pharma and Diagnostics, Mannheim, Germany) was mounted onto their shaved back. Background signal was measured for 5 minutes before injection of 5 mg/100g body weight of FITC-sinistrin. Transcutaneous fluorescence was then continuously measured for one hour in conscious freely moving animals. mGFR (µL/min) was calculated from the decrease in fluorescence intensity over time (i.e. plasma half-life of FITC-sinistrin). A detailed presentation of the data output and GFR calculation from signal decay is available in a recent publication.[S2]

Blood pressure telemetry: 12–16-week-old POD-ATTAC mice were housed in individual cages and implanted with PA-C10 radiotelemetry devices (Data Sciences International, DSI, USA) measuring heart rate and systolic and diastolic blood pressure, as described previously.[S3] Surgery was performed under isoflurane anaesthesia and buprenorphine was used for perioperative pain management. Mice were allowed to recover for 12 days after surgery and were housed under a 12-hour dark/light cycle (lights off at 19:00, on at 07:00) with free access to food and water. Each blood pressure recording session lasted 24 hours (09:00–09:00). Blood pressure measurements were collected using Ponemah software (DSI), recording for 15 seconds every minute. The first 60 minutes of each recording were excluded to eliminate the blood pressure spike associated with stress at the start of the experiment. Averages were calculated for the full, dark, and light periods. A baseline recording was performed before introduction of the experimental 0.8K or 1.3K dietary bolus, followed by a second recording after one week on diet. Podocyte apoptosis was then induced by five daily injections of AP20187 as described above. A third recording was performed 10 days after the first injection to study the acute effect on blood pressure, and a fourth 28 days after the first injection to study the long-term effect.

For validation of the MRA class effect in the POD-ATTAC model, spironolactone was added to the dietary bolus at 250 mg/kg/day from the third day after the first AP20187 injection until the end of the experiment.

*Plasma and urine biochemistry*

Plasma potassium and sodium were measured by potentiometry using the Epoc® blood analysis system (Siemens Healthcare, Erlangen, Germany). Retro-orbital venous blood was sampled under terminal anaesthesia with ketamine and xylazine immediately prior to exsanguination and analysed immediately. The remainder of the blood was collected in heparinised plasma tubes and centrifuged. Plasma was stored at −80°C. Aldosterone was measured from plasma samples by radioimmunoassay (R-CW 100, Diasource) at the University Hospital of Lausanne, Switzerland.

For measurement of urinary potassium, mice were placed on their assigned potassium diet for one week, then acclimatised to Tecniplast metabolic cages for 48 hours; urine output was collected over the subsequent 24-hour period. The assigned dietary potassium content was maintained throughout the metabolic cage period. Urinary potassium was measured by flame photometry (IL943; Instrumentation Laboratory, Ramsey, MN, USA) at the Centre de Recherche des Cordeliers, Sorbonne Université, Université de Paris, Paris, France.[S4]

*Histology*

Paraformaldehyde-fixed samples were washed and embedded in paraffin. Five-micrometre sections were deparaffinised and stained with Fast Green/Sirius red.[S5,S6] Whole slides were scanned with a digital slide scanner Axioscan 7 (Carl Zeiss, Oberkochen, Germany) under white and circularly polarised light, and analysed in QuPath version 0.6.0.[S7] A custom pixel classifier, trained on representative images with dedicated annotations, was used to detect collagen staining and polarisation signal, an example is provided in Ssupplementary Figure 11.

Immunofluorescence microscopy was performed on 5 μm deparaffinised sections from paraformaldehyde-fixed samples. Primary antibodies were incubated overnight at 4°C; secondary antibodies were incubated at room temperature for 1 hour, followed by a brief DAPI incubation. Whole slides were scanned with the same scanner and analysed in QuPath. The InstanSeg plugin was used to segment cell objects from DAPI signal.[S8] A random tree object classifier was then trained on representative images from each cohort to identify cells expressing the marker of interest. An example of classifier output is provided in Supplementary Figure 12 for the POD-ATTAC model, in which non-specific proteinuric cast signal was excluded from PDGFRβ quantification. An example for αSMA object classification is also provided in Supplementary Figure 13. Negative controls were performed in the absence of primary or secondary antibody (data not shown).

PDGFRβ was detected following antigen retrieval in Tris-EDTA buffer using a mouse monoclonal anti-PDGFRβ primary antibody (Abcam, ab69506; 1:100), followed by an Alexa Fluor 488-conjugated goat anti-mouse secondary antibody (ThermoFisher Scientifice, A-11029, 1:100). αSMA was detected following antigen retrieval in citrate buffer using a homemade mouse monoclonal anti-αSMA primary antibody (University of Geneva antibody platform, kindly provided by Pr M-L Bochaton-Piallat; 1:100),[S9,S10] followed by an Alexa Fluor 594-conjugated goat anti-mouse secondary antibody (Jackson ImmunoResearch, 115-585-206; 1:100).

*Western blotting*

Kidney tissue samples were homogenised in 200 µL of lysis buffer (imidazole 25.2 mM; EDTA 1 mM; sucrose 300 mM, NP-40 0.5% v/v; sodium fluoride 50 mM; $Na_3VO_4$ 2 mM; protease inhibitors). Protein concentrations were determined using the bicinchoninic acid protein assay (Thermo Scientific, Pierce). Proteins were separated by SDS-PAGE and transferred onto polyvinylidene difluoride membranes (Immobilon-P, Millipore). Membranes were blocked for 1 hour in 5% milk in TBSt buffer and incubated with primary antibodies diluted in 5% milk/TBSt overnight at 4°C, followed by HRP-conjugated secondary antibodies for 1 hour at room temperature. Enhanced chemiluminescence (WesternBright ECL, Advansta, San Jose, CA, USA) was used for detection. Protein levels were quantified using ImageJ version 1.54g.[S11]

Results are expressed as the ratio of the densitometry of the band of interest to the loading control, normalised to the average of the reference group. Fibronectin was detected using a rabbit polyclonal anti-fibronectin antibody (Abcam, ab2413; 1:1000) and E-cadherin using a mouse monoclonal anti-E-cadherin antibody (BD Biosciences, cat. no. 610181; 1:2000), with HRP-conjugated goat anti-rabbit (BD Biosciences, cat. no. 554021; 1:5000) and goat anti-mouse (BD Biosciences, cat. no. 554002; 1:5000) secondary antibodies respectively.

*RNA extraction and real-time qPCR*

Total RNA was extracted from kidney tissue using the NucleoSpin RNA II kit (Macherey-Nagel) according to the manufacturer's instructions. RNA concentration and purity were assessed by ultraviolet spectrophotometry. One microgram of RNA was used to synthesise cDNA using the qScript cDNA Supermix (Quanta Biosciences). Real-time PCR was performed on 10 ng of cDNA diluted 1:6 (v/v) using 0.5 µM of each primer and 6 µL of PowerUp SYBR Green Master Mix (Applied Biosystems) in a final reaction volume of 12 µL. Duplicate amplification reactions were performed on a QuantStudio 5 real-time PCR system (Thermo Fisher Scientific). Mouse acidic ribosomal phosphoprotein p0 (Rplp0) was used as the internal reference standard. Nr3c2 deletion in MR-KO$_{tub}$ mice was confirmed by qPCR on cDNA from contralateral kidney tissue using the following primers listed in Supplementary Table 2.

*gDNA PCR*

Routine PCR on genomic DNA from phalanges was used to genotype transgenic animals, with primers listed in Supplementary Table 2. To confirm recombination of *Nr3c2* in MR-KO$_{msc}$

animals, PCR was performed on ear pinnae collected at sacrifice. Ear pinnae were stored at 4°C and lysed in 25 mM NaOH, 0.2 mM EDTA for 30 minutes at 95°C. One microlitre of lysate was amplified with 1 µM of each primer and 10 µL of Phire Green Hot Start II Master Mix (Applied Biosystems) in a final volume of 20 µL. Samples were run on a 3% agarose gel in Tris-acetate-EDTA buffer. Primers were designed in the intronic regions flanking Nr3c2 exon 3, outside the loxP sites, to produce a shorter amplicon when recombination had occurred: forward, 5'-GTTTACGGCGTGGTGATCTT-3'; reverse, 5'-ATATTCCCGAACAGCGCATA-3'.

S2. Chassot A, Feraille E, Sassi A. Claudin-3 is expressed at the macula densa and is associated with altered glomerular filtration in claudin-3 knockout mice. *Front Physiol* [ahead of print].

S3. Wang Y, Thatcher SE, Cassis LA. Blood pressure monitoring using radio telemetry method in mice. In: Thatcher SE, editor. *The Renin-Angiotensin-Aldosterone System: Methods and Protocols*. New York: Springer; 2017. p. 75–85.

S4. Wolf MJ, El Moghrabi S, Palacios-Ramirez R, et al. The novel mineralocorticoid receptor modulator balcinrenone protects against diet-induced cardiac microvascular dysfunction and plasma potassium elevation in mouse models. *PLoS One* 2026;21(2):e0341078.

S5. Junqueira LCU, Bignolas G, Brentani RR. Picrosirius staining plus polarization microscopy, a specific method for collagen detection in tissue sections. Histochem J 1979;11(4):447–55.

S6. Street JM, Souza ACP, Alvarez-Prats A, et al. Automated quantification of renal fibrosis with Sirius Red and polarization contrast microscopy. Physiol Rep 2014;2(7):e12088.

S7. Bankhead P, Loughrey MB, Fernández JA, et al. QuPath: Open source software for digital pathology image analysis. *Sci Rep* 2017;7(1):16878.

S8. Goldsborough T, et al. A novel channel invariant architecture for the segmentation of cells and nuclei in multiplexed images using InstanSeg. *bioRxiv* 2024. doi:10.1101/2024.09.04.611150.

S9. Skalli O, Ropraz P, Trzeciak A, Benzonana G, Gillessen D, Gabbiani G. A monoclonal antibody against alpha-smooth muscle actin: a new probe for smooth muscle differentiation. *J Cell Biol* 1986;103(6 Pt 2):2787–96.

S10. Darby I, Skalli O, Gabbiani G. Alpha-smooth muscle actin is transiently expressed by myofibroblasts during experimental wound healing. *Lab Invest* 1990;63(1):21–9.

S11. Schneider CA, Rasband WS, Eliceiri KW. NIH Image to ImageJ: 25 years of image analysis. *Nat Methods* 2012;9(7):671–5.

Supplementary Table 1 : Sodium and Potassium and anionic content of all diets*

| **Sodium content (from sodium chloride)** | **Potassium content** | **Accompanying anion to potassium** | **Naming convention** |
|---|---|---|---|
| 0.32% (normal sodium) | 0.01% (low potassium) | Chloride | 0.01K |
| 0.32% (normal sodium) | 0.8% (normal potassium) | Chloride | 0.8K |
| 0.32% (normal sodium) | 1.3% (medium high potassium) | Chloride | 1.3K |
| 0.32% (normal sodium) | 2% (high potassium) | Chloride | 2K |
| 0.32% (normal sodium) | 0.8% (normal potassium) | Citrate | 0.8K-Cit |
| 0.32% (normal sodium) | 1.3% (medium high potassium) | Citrate | 1.3K-Cit |
| 0.32% (normal sodium) | 2% (high potassium) | Chloride and Citrate† | 2K-ClCit |
| 1.26% (high sodium) | 0.01% (low potassium) | Chloride | HS 0.01K |
| 1.26% (high sodium) | 0.8% (normal potassium) | Chloride | HS 0.8K |
| 1.26% (high sodium) | 1.3% (medium high potassium) | Chloride | HS 1.3K |
| 1.26% (high sodium) | 2% (high potassium) | Chloride | HS 2K |

*Dietary boluses were prepared from a custom purified powdered diet base (Granovit AG; full details in Methods). Ion concentrations are expressed as percentage weight/weight of the final bolus and refer to the elemental content rather than the complexed salt. Not all combinations were used in every model; diet assignments per experiment are specified in the relevant figure legends and results sections †2K-ClCit contained 0.8% K from potassium chloride and 1.2% K from potassium citrate.

Supplementary Table 2: List of primers used for PCR and qPCR

| Gene/Transgene | Primer | Sequence 5' - 3 ' |
|---|---|---|
| **Genotyping , gDNA PCR)** | | |
| $MR^{lox/lox}$ | Forward | CTCGAGATCTGAACTCCAGGC |
| | Reverse 1 | CCTAGAGTTCCTGAGCTGCTGA |
| | Reverse 2 | TAGAAACACTTCGTAAAGTAGAGCT |
| Pax8-rtTA | Forward | CCATGTCTAGACTGGACAAGA |
| | Reverse | CTCCAGGCCACATATGATTAG |
| Cre-LC1 (Genotyping) | Forward | TCGCTGGATTACCGGTCGATG |
| | Reverse | CCATGAGTGAACGAACCTGGTCG |
| PDGFRβ-CreERT2 (Genotyping) | Forward | CCACCTTGAATGAAGTCAACAC |
| | Reverse 1 | ACATGTCCATCAGGTTCTTGC |
| | Reverse 2 | AGCTTGTGGCAGTGTAGCTG |
| $Lyz2^{tm1(cre)Ifo}$ (Genotyping) | Forward | CTTGGGCTGCCAGAATTTCTC |
| | Reverse 1 | CCCAGAAATGCCAGATTACG |
| | Reverse 2 | TTACAGTCGGCCAGGCTGAC |
| POD-ATTAC (Genotyping) | Forward | GAAAGTGCCCAAACTTCAGAGCATTAGG |
| | Reverse | AACTGAGATGTCAGCTCATAGATGGGGG |
| **qPCR Gene Expression** | | |
| Nr3c2 (Gene expression) | Forward | ATCTGTGTGCTGGAAGAAATGAC |
| | Reverse | TTTAACTTCCCCAGCTTCTTTG |
| Col1α1 (Gene expression) | Forward | GGCAAGAATGGAGATGATGG |
| | Reverse | ACCATCCAAACCACTGAAGC |
| PDGFRβ | Forward | TGTGATCGAGAATGGCTACG |
| | Reverse | ACAGAAGGCATCGGATAAGC |
| αSMA | Forward | TCAGCGCCTCCAGTTCCT |
| | Reverse | AAAAAAAACCACGAGTAACAAATCAA |

a

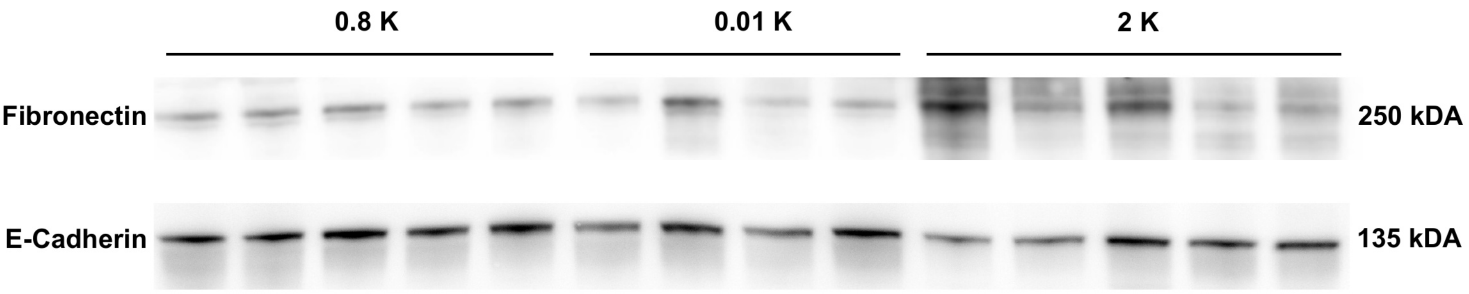


b

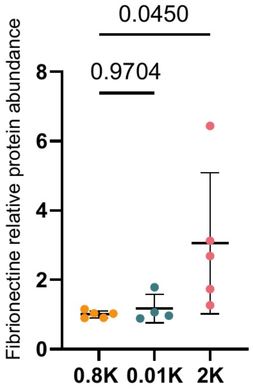


**Supplementary Figure 1.** Evaluation of renal fibrosis after unilateral ureteral obstruction (UUO). **a**: Representative Western blots for fibronectin and E-cadherin in obstructed kidneys from animals fed 0.8K (n=5), 0.01K (n=4), or 2K (n=5) diets. **b**: Fibronectin band intensity normalized to E-cadherin and expressed relative to the 0.8K group mean (one-way ANOVA, p=0.0484

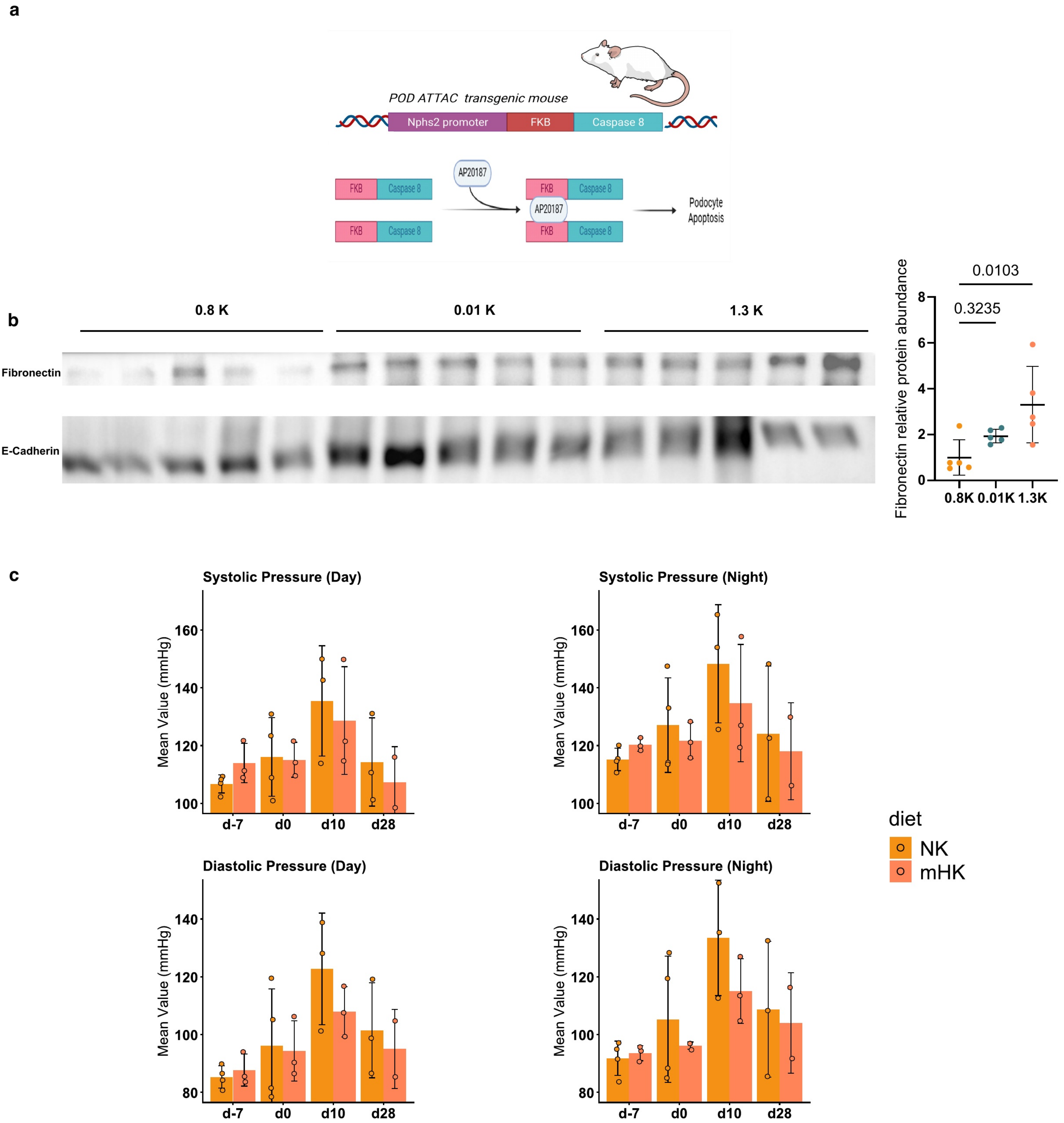


**Supplementary Figure 2.** Evaluation of renal fibrosis after podocyte apoptosis induction (POD ATTAC). **a** The POD ATTAC model is a transgenic mouse line expressing a fusion protein of an FKBP fragment and Caspase-8 specifically in podocytes. Injection of AP20187 induces dimerization of Caspase-8 via its FK506 binding (FKBP) domains, triggering podocyte apoptosis. **b** Left: Representative Western blots for fibronectin and E-cadherin in kidneys from POD ATTAC mice fed 0.8K (n=5), 0.01K (n=5), or 1.3K (n=5) diets. Right: Fibronectin band intensity normalized to E-cadherin and expressed relative to the 0.8K group mean (one-way ANOVA, p=0.0176). **c** Blood pressure was recorded over 24 hours in a subset of mice implanted with PA-C10 telemetry devices at four time points: before dietary intervention (day −7), after one week on diet and immediately prior to the first injection (day 0), during the acute injury phase (day 10), and during the chronic phase (day 28). Mean day and night period systolic and diastolic blood pressure is shown for the 0.8K and 1.3K groups.

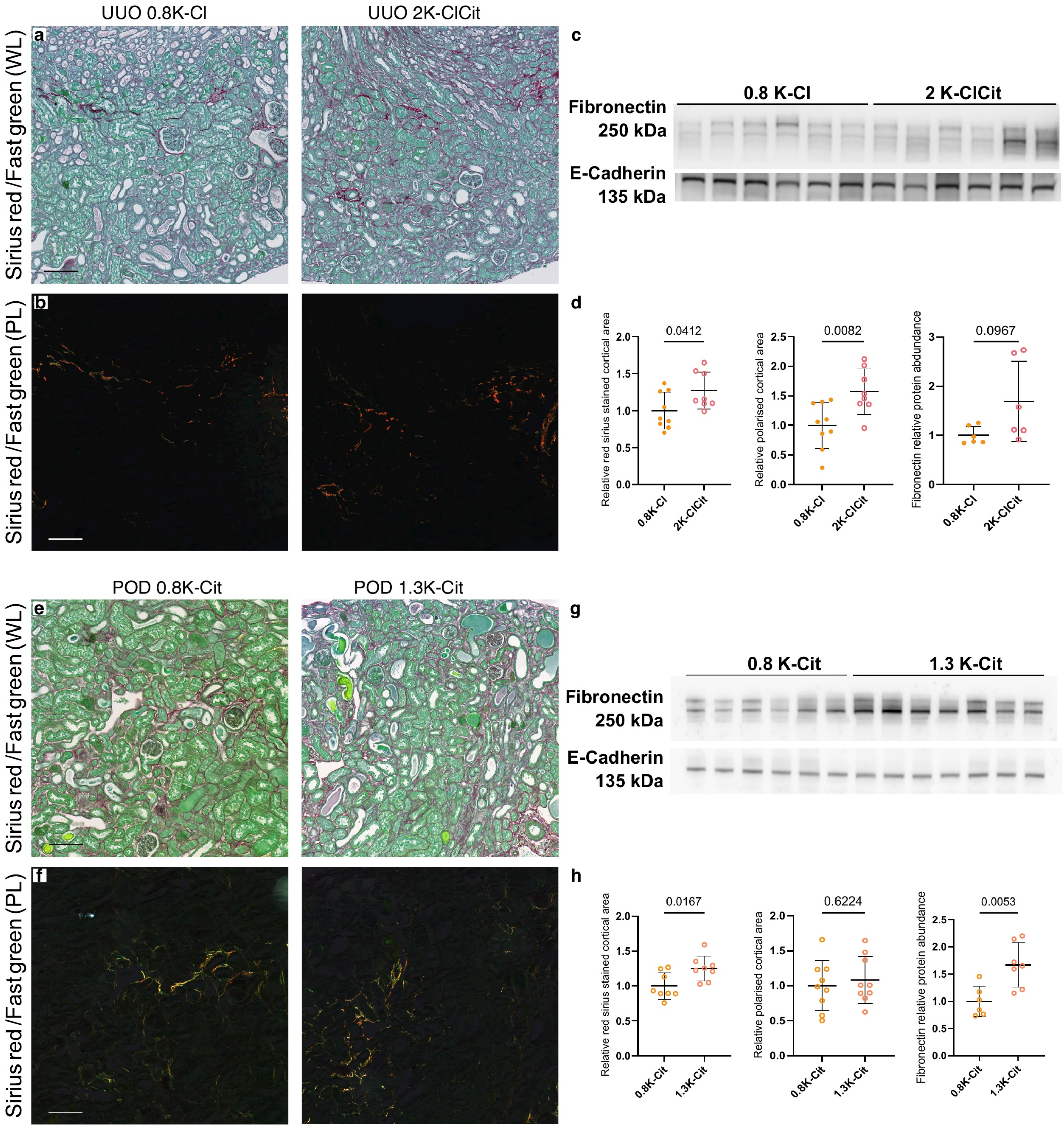


**Supplementary figure 3** . Effect of anion type on the potassium–fibrosis relationship. We studied the effect of potassium citrate either added to a baseline potassium chloride diet (UUO) or as the sole source of potassium (POD-ATTAC). Fibrosis was induced by UUO under the reference 0.8K-Cl potassium chloride diet or a 2K-ClCit (0.8K from KCl and 1.2 from K-citrate) diet (**a, b, c, d**) or podocyte apoptosis under diet strictly containing potassium citrate, either 0.8K-Cit or 1.3K-Cit (**e, f, g, h**). **a, b, e, f** Representative coronal paraffin sections stained with Sirius red under white light (WL; **a, e**) and polarized light (PL; **b, f**); scale bar 100 µm. c Representative Western blots for fibronectin and E-cadherin for UUO mice under these diets. . **d** Left: Quantification of cortical Sirius red area (WL) from a, normalized to the 0.8K-Cl mean (n=9/8). Middle: Quantification of cortical Sirius red polarized area from b, normalized to the 0.8K-Cl mean (n=9/8). Right: Fibronectin band intensity normalized to E-cadherin from c and expressed relative to the 0.8K-Cl group mean (n=6/6). **g** Representative Western blots for fibronectin and E-cadherin for POD ATTAC mice under citrate diets. **h** Left: Quantification of cortical Sirius red area (WL) from e, normalized to the 0.8K-Cit mean (n=8/8). Middle: Quantification of cortical Sirius red polarized area from f, normalized to the 0.8K-Cit mean (n=8/8). Right: Fibronectin band intensity normalized to E-cadherin from g and expressed relative to the 0.8K-Cit group mean (n=6/7).

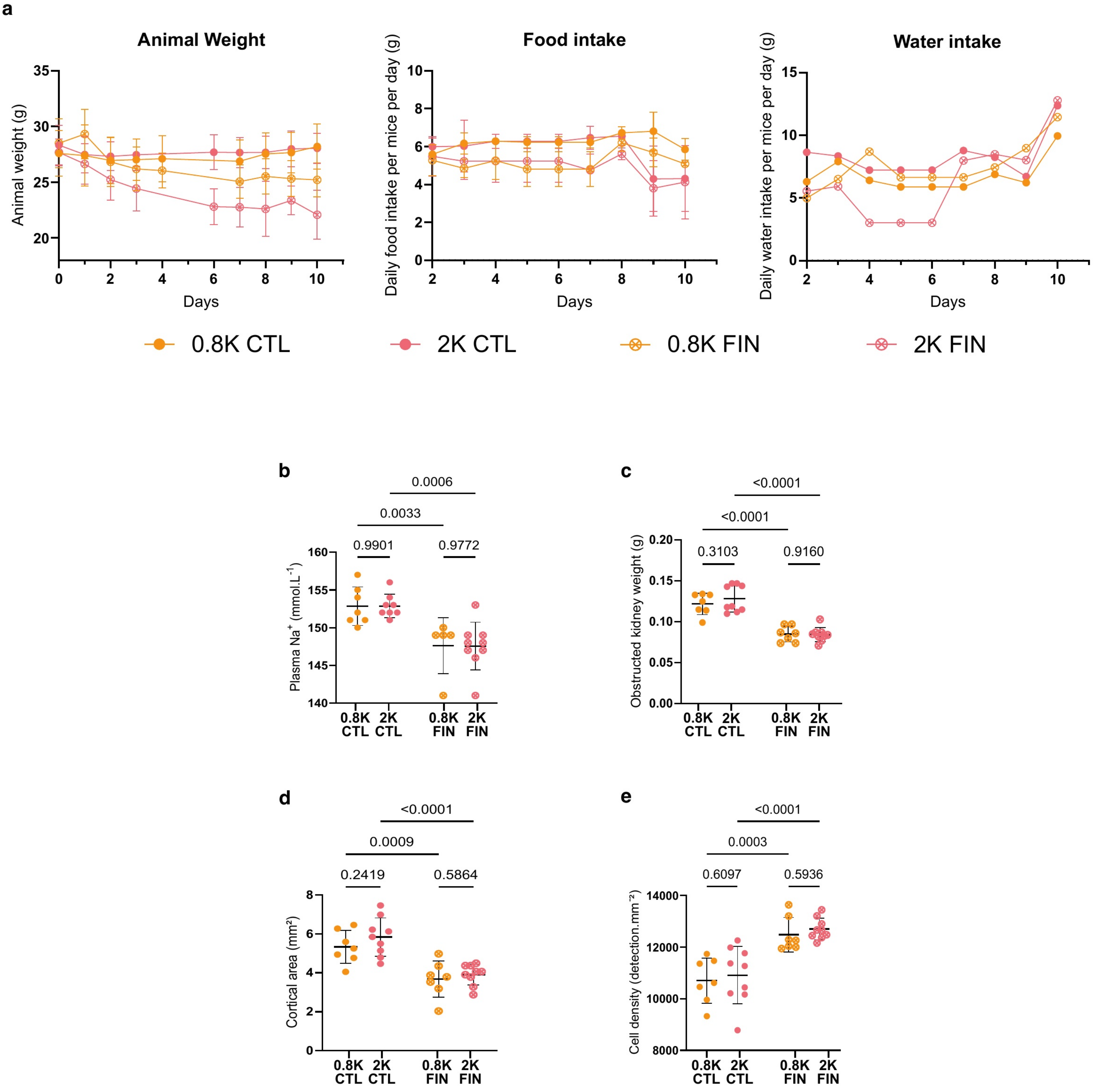


**Supplementary Figure 4.** Physiological effects of combined finerenone treatment and dietary potassium. (n unless stated otherwise: 0.8K CTL = 7, 2K CTL = 9, 0.8K FIN = 7, 2K FIN = 9) **a** Longitudinal measurements over the 10-day experiment. Left: Body weight (g), recorded on workdays only. Middle: Average daily food intake per mouse (g), estimated from cage food weight. Right: Average daily water intake per mouse (g), estimated from bottle weight. Physiological measurements at sacrifice: **b** Plasma sodium (n reduced due to assay failure in some samples, n= 7/8/5/9, same group order as above) (potentiometry; two-way ANOVA: p interaction=0.9766, p treatment <0.0001, p diet=0.9900). **c** Obstructed kidney weight (g; two-way ANOVA: p interaction=0.5623, p treatment <0.0001, p diet=0.5175). **d** Cortical area measured on Sirius red slides (two-way ANOVA: p interaction=0.6517, p treatment <0.0001, p diet=0.2272). **e** Cell density in cortex from αSMA immunofluorescence, expressed as cell count per cortical surface area (two-way ANOVA: p interaction=0.9868, p treatment <0.0001, p diet=0.4614).

a

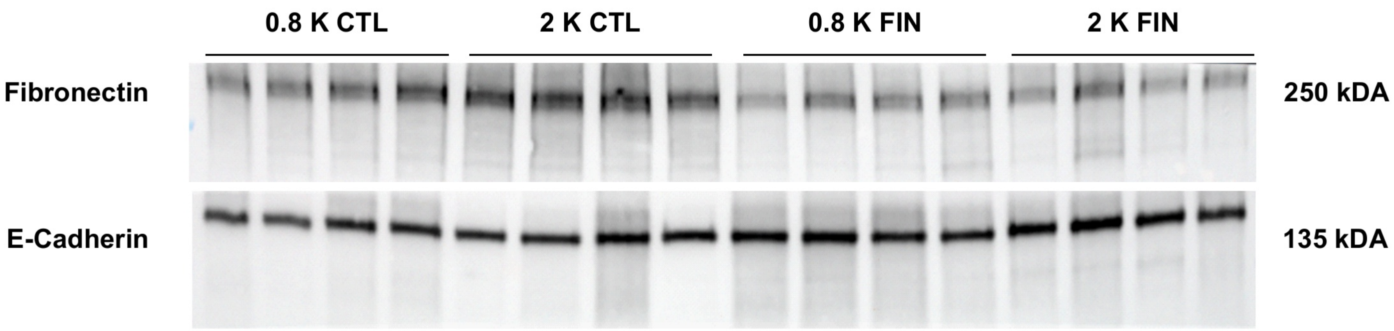


b

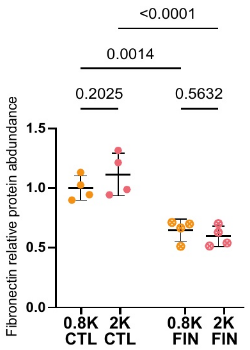


**Supplementary Figure 5.** Finerenone reduces high-potassium-induced fibrosis. **a**: Representative Western blots for fibronectin and E-cadherin in kidneys from the 0.8K untreated, (CTL),2K CTL, 0.8K Finerenone treated (FIN) and 2K FIN groups, n=4 per group. **b**: Fibronectin band intensity normalized to E-cadherin and expressed relative to the 0.8K control group mean (n= 4 for each group ,two-way ANOVA: p interaction=0.6848, p treatment=0.0004, p diet=0.0767)

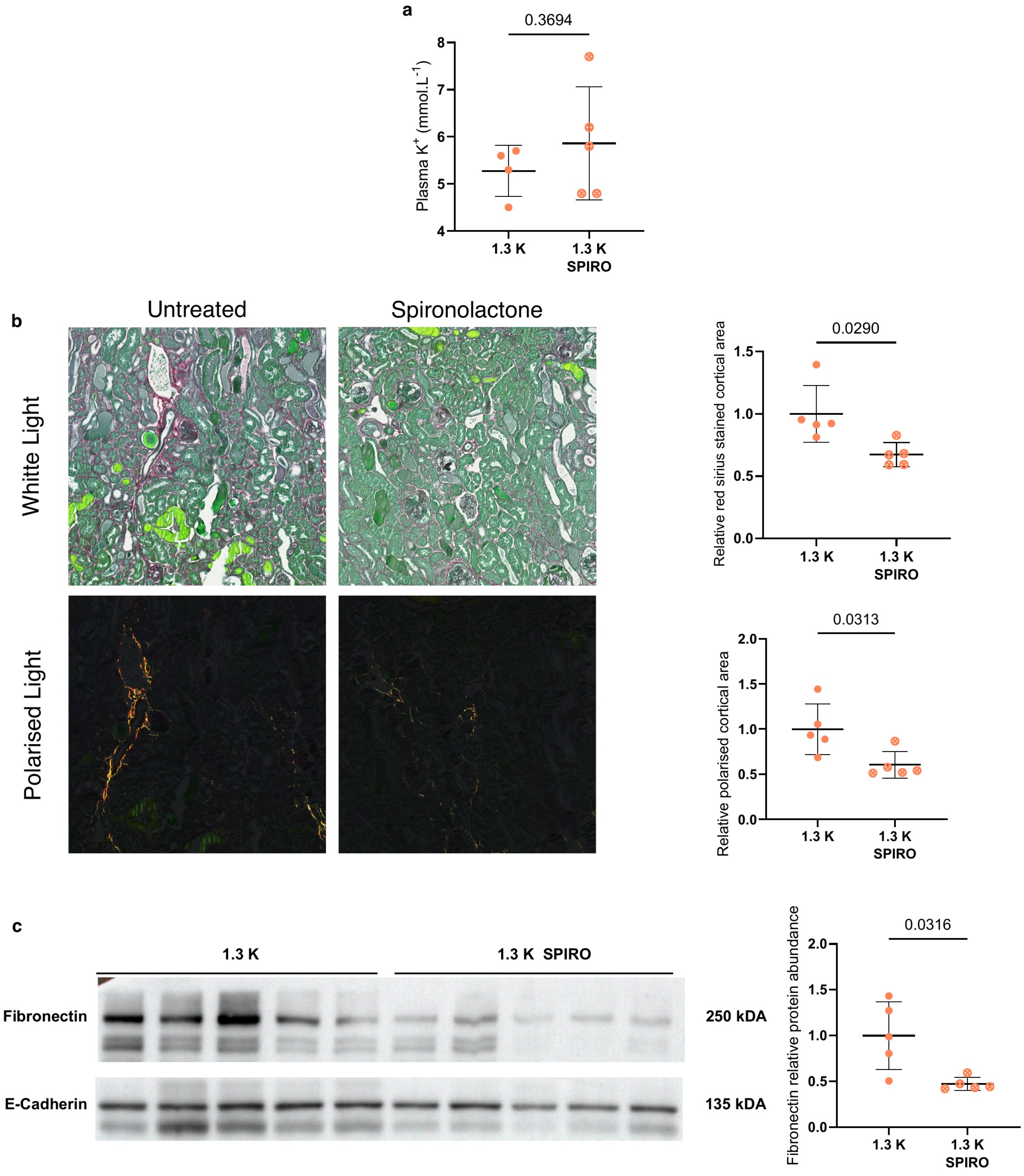


**Supplementary Figure 6**. Effect of spironolactone on renal fibrosis in high-potassium-fed mice. POD ATTAC mice were placed on a medium-high (1.3K) potassium diet on the day of the first of five daily injections to induce podocyte apoptosis, with (SPIRO) or without (Untreated) spironolactone added to the diet from day one, and sacrificed 20 days later for biochemical and histological analysis.(n = 5/group; 1.3K in panel a: n = 4, one measurement failed, all comparisons by two tailed Welch's t-test) **a** Plasma potassium at sacrifice (potentiometry). **b** Representative coronal paraffin sections stained with Sirius red under white light (WL, top) and polarized light (PL, bottom). Right: Quantification of cortical Sirius red area — WL (top) and PL (bottom) — normalized to the 1.3K mean. **c** *Left*: Representative Western blots for fibronectin and E-cadherin in kidneys from mice fed 1.3K or 1.3K SPIRO. *Right*: Fibronectin band intensity normalized to E-cadherin and expressed relative to the 1.3K group mean.

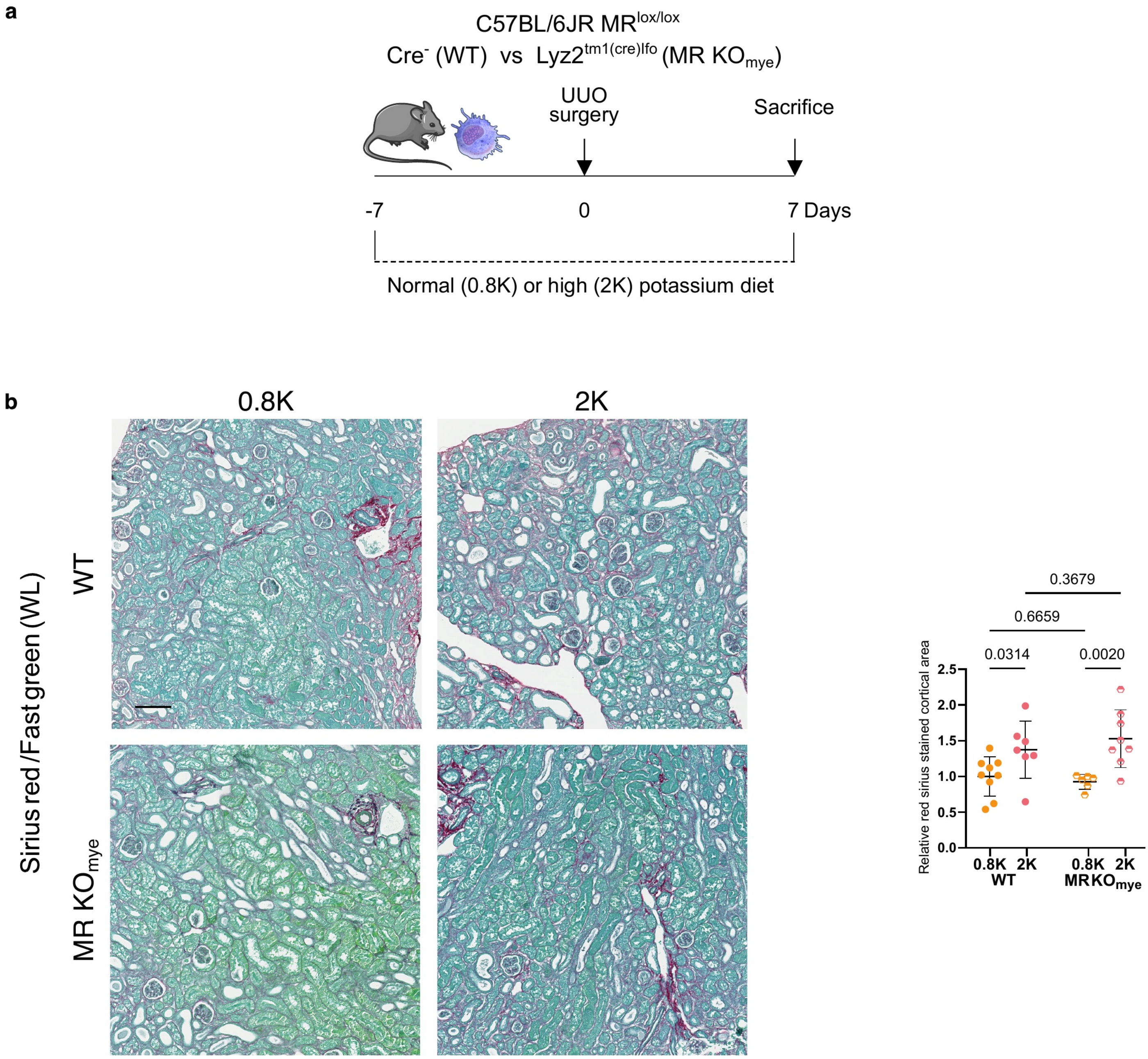


**Supplementary figure 7.** Effect of myeloid mineralocorticoid receptor knockout on renal fibrosis. **a** $MR^{lox/lox}$ $Lyz2^{tm1(cre)Ifo}$ mice, in which Cre is constitutively expressed in myeloid cells ($MR\text{-}KO_{mye}$), and their $MR^{lox/lox}Cre^{-}$ littermates (WT) were placed on 0.8K or 2K diets one week before UUO and sacrificed 7 days post-procedure ( n= 0.8K WT:9 ; 2K WT:7 ; 0.8K MR-KOmye: 6 ; 2K MR-KOmye: 8) . **b** Representative coronal paraffin sections stained with Sirius red under white light; scale bar 100 µm and quantification of cortical Sirius red area, normalised to the 0.8K WT mean (two-way ANOVA: p interaction=0.3490, p genotype=0.7438, p diet=0.0004).

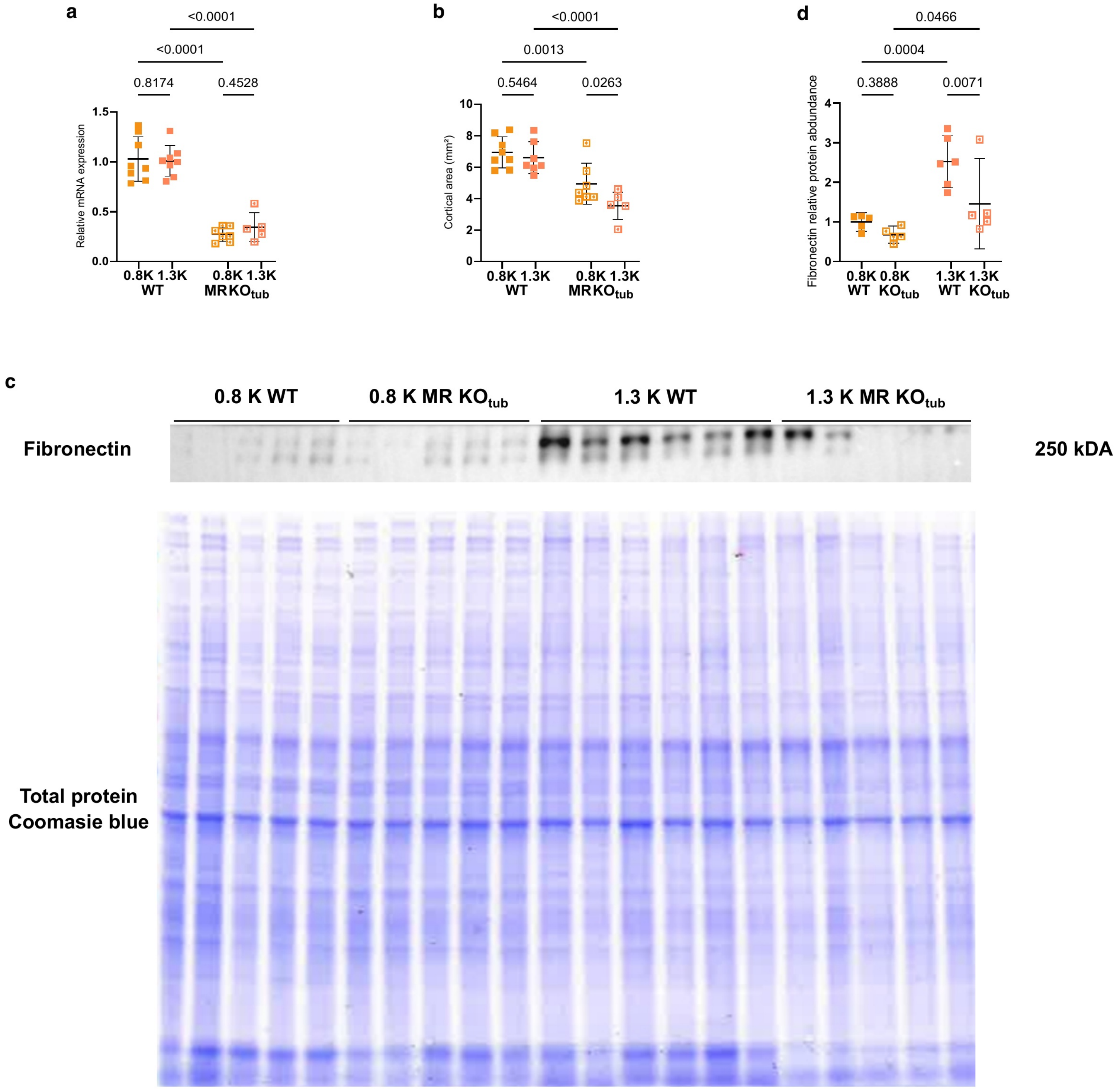


**Supplementary Figure 8.** Effect of tubular mineralocorticoid receptor knockout on renal fibrosis. ( for a and b n= 0.8K WT: 8 ; 2K WT: 7 ; 0.8K MR-KOtub : 7 ; 2K MR-KOtub : 5) **a** Relative *Nr3c2* mRNA expression from contralateral cortical kidney tissue (two-way ANOVA: p interaction=0.4713, p genotype<0.0001, p diet=0.6719). **b** Cortical section area from obstructed kidneys (mm²; two-way ANOVA: p interaction=0.2235, p genotype<0.0001, p diet=0.0525). **c** Representative Western blots for fibronectin in kidneys from all four groups. Loading was normalized to total protein (Coomassie) rather than E-cadherin due to antibody unavailability at the time of blotting.. Lanes were loaded in the order 0.8K WT (n=5), 0.8K MR-KOtub (n=5), 2K WT (n=6), 2K MR-KOtub (n=5), not 0.8K WT, 2K WT, 0.8K KO, 2K KO as in other panels. **d** Fibronectin band intensity normalised to total protein and expressed relative to the 0.8K WT group mean (two-way ANOVA: p interaction=0.1580, p genotype=0.0135, p diet=0.0003).

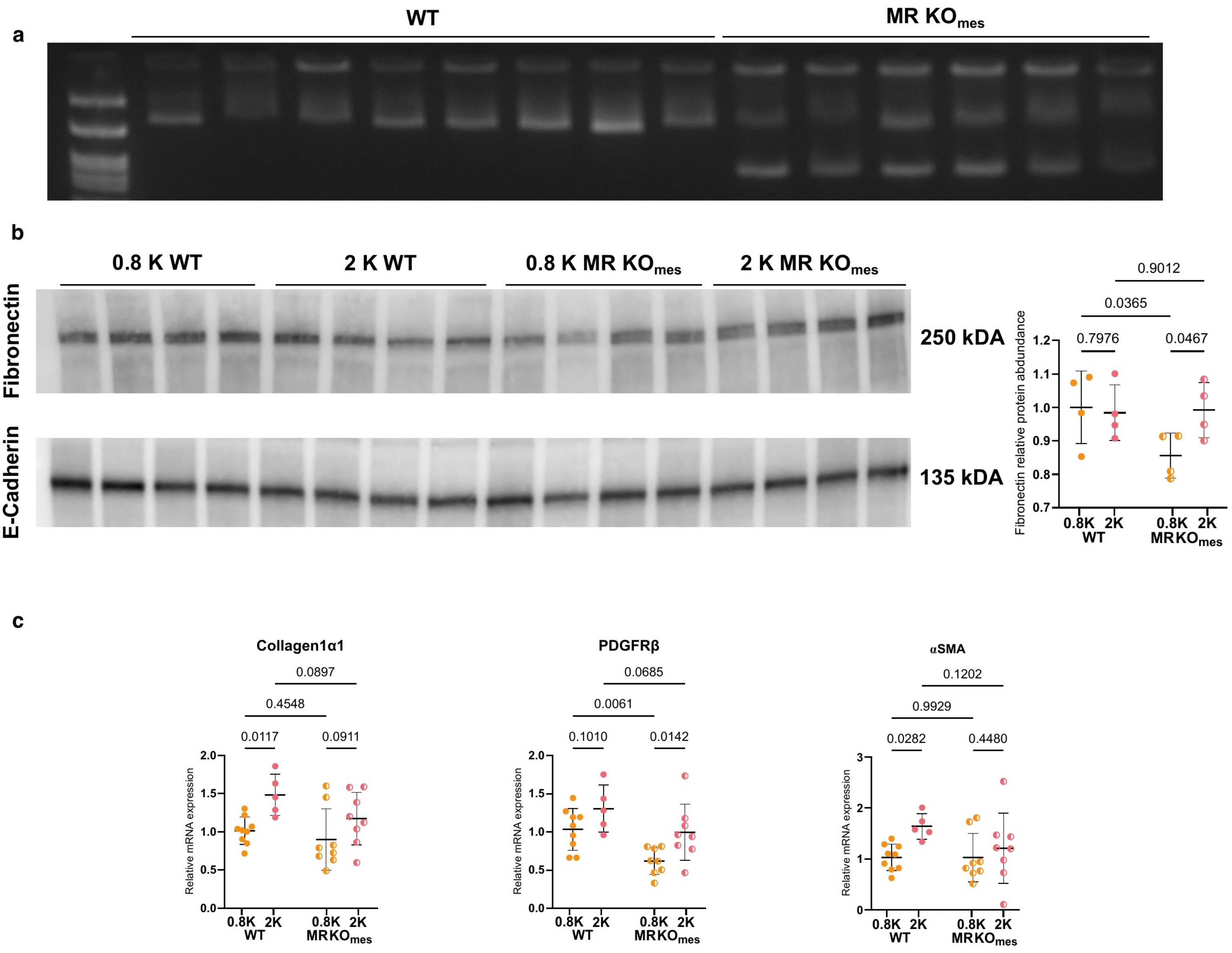


**Supplementary Figure 9.** Effect of mesenchymal mineralocorticoid receptor knockout on renal fibrosis. **a** Representative PCR gel from fibroblast-rich ear pinnae. Primers flanking *Nr3c2* exon 3 in intronic regions outside the loxP sites amplify a band of size dependent on recombination status. The 8 WT animals show a single longer band, while the 6 MR-KO$_{mes}$ animals display both bands, confirming successful recombination. **b** Left: Representative Western blots for fibronectin and E-cadherin in kidneys cortex extracts (n= 0.8KWT: 4 ; 2K WT: 4 ; 0.8K MR-KO$_{mes}$ : 4, 2K MR-KO$_{mes}$: 4). Right: Fibronectin band intensity normalized to E-cadherin and expressed relative to the 0.8K WT group mean. Results did not reach statistical significance in this cohort (two-way ANOVA: p interaction=0.1050, p genotype=0.1415, p diet=0.1921) ), possibly reflecting technical limitations associated with sample number constraints and gel format. **c** Relative mRNA expression after UUO (n= 0.8KWT: 9 ; 2K WT: 5 ; 0.8K MR-KO$_{mes}$ : 9, 2K MR-KO$_{mes}$: 9) of Collagen1α1 (two-way ANOVA: p interaction=0.4036, p genotype=0.0782, p diet=0.0037), PDGFRβ (p interaction=0.6297, p genotype=0.0023, p diet=0.0056), αSMA (p interaction=0.2346, p genotype=0.2302, p diet=0.0335).

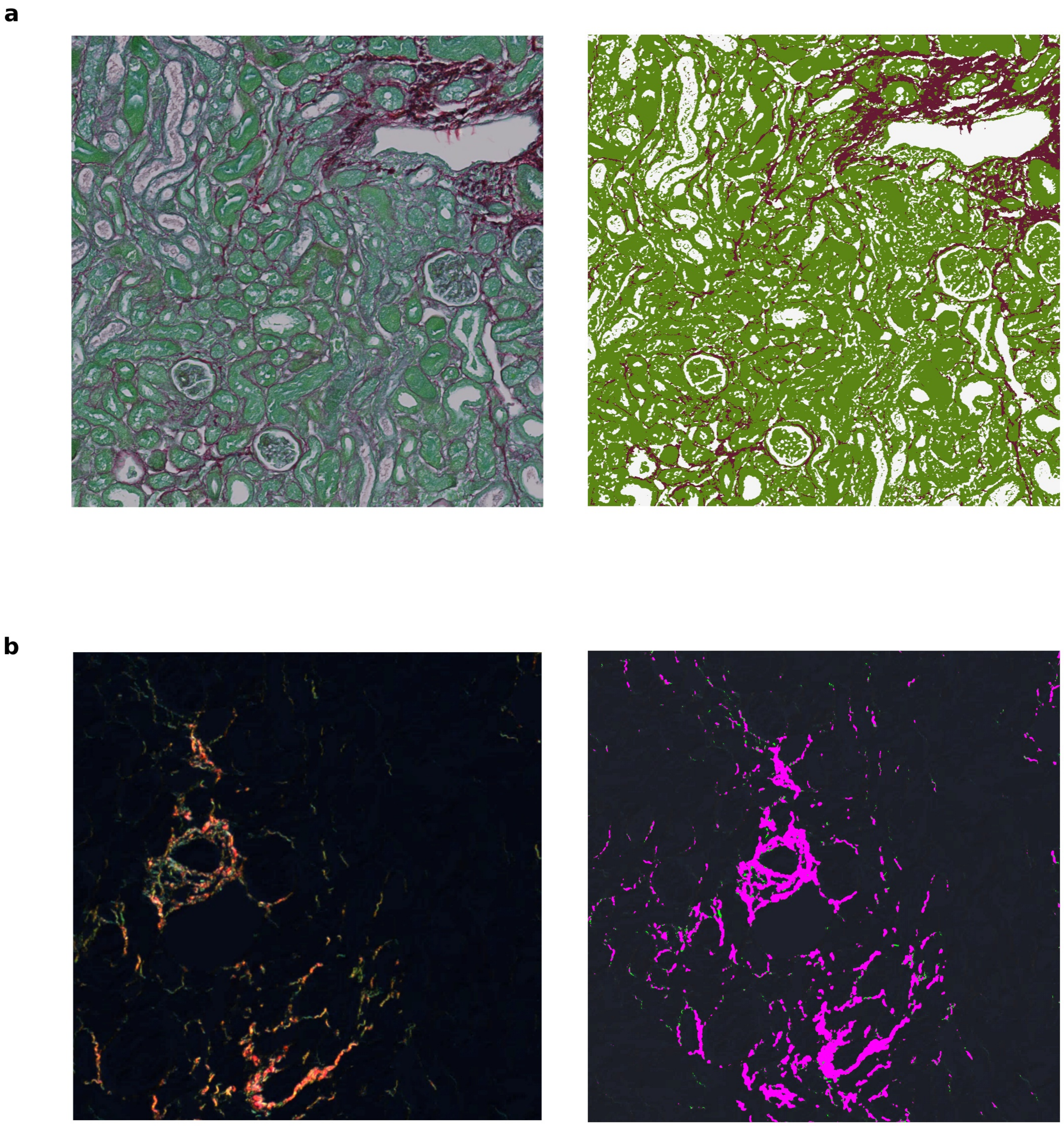


**Supplementary Figure 10.** Example of pixel classification for Sirius red/Fast Green staining under white and polarised light. Representative images from an obstructed kidney stained with Sirius red/Fast Green, observed under white light (a) and polarised light (b). In each panel, the left image shows the native acquisition and the right image shows the corresponding pixel classification output. For white light (a), pixels are classified as either negative (unstained), counterstained (green) or positive (red) for Sirius red staining. For polarised light (b), green and red birefringence signals were detected first in separate channels and summed for classification.

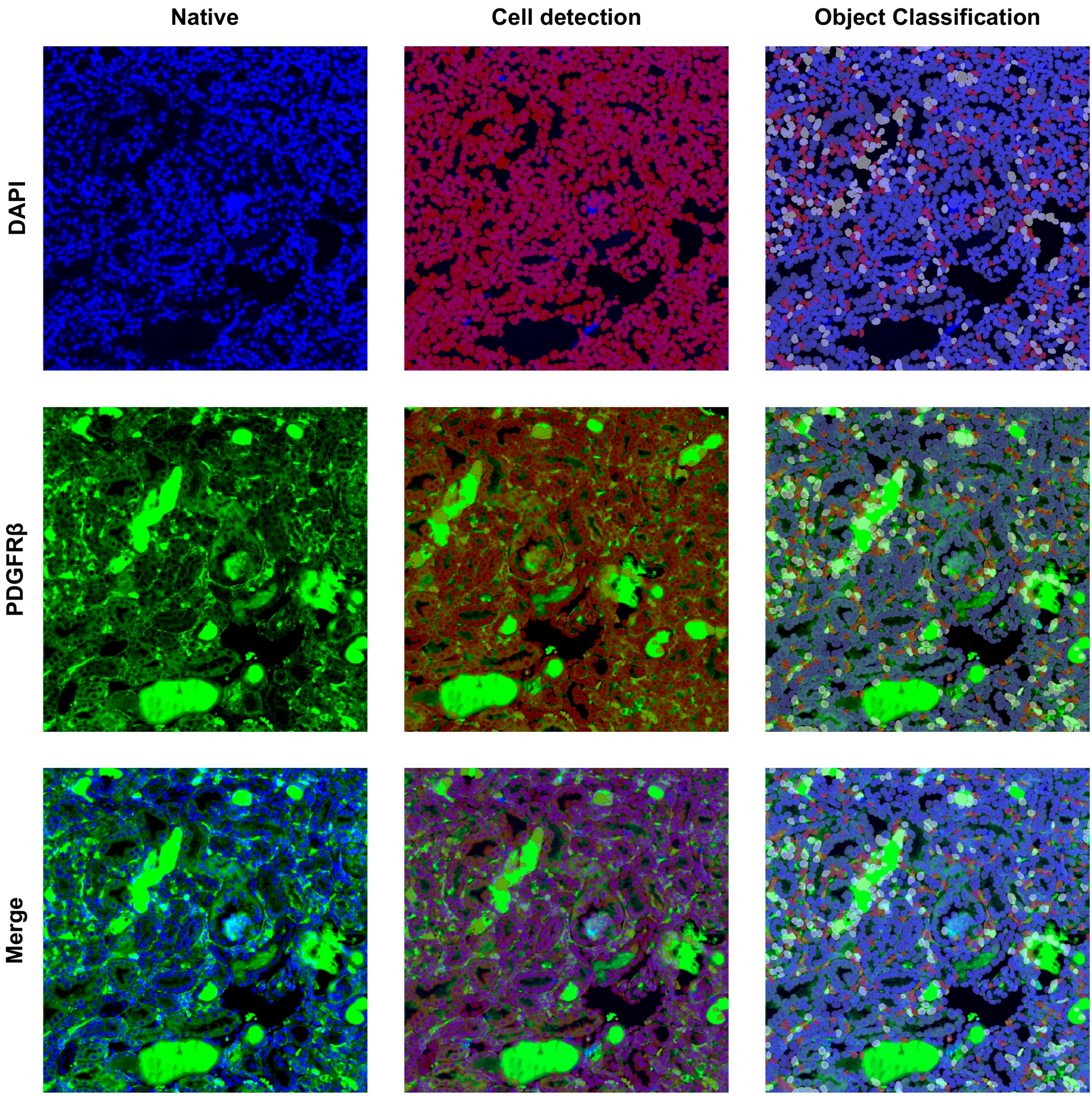


**Supplementary Figure 11.** Example of PDGFRβ-positive interstitial cell detection. Representative images from a POD-ATTAC cohort slide immunostained with DAPI and anti-PDGFRβ antibody, illustrating the object classification workflow. Rows show, from top to bottom: DAPI, PDGFRβ, and merged channels. Columns show, from left to right: native image, cell detection overlay, and object classification output. Blue denotes PDGFRβ-negative cells; white denotes PDGFRβ-positive cells excluded from quantification (adjacent fluorescence signal from tubular casts); red denotes PDGFRβ-positive interstitial cells retained for quantification.

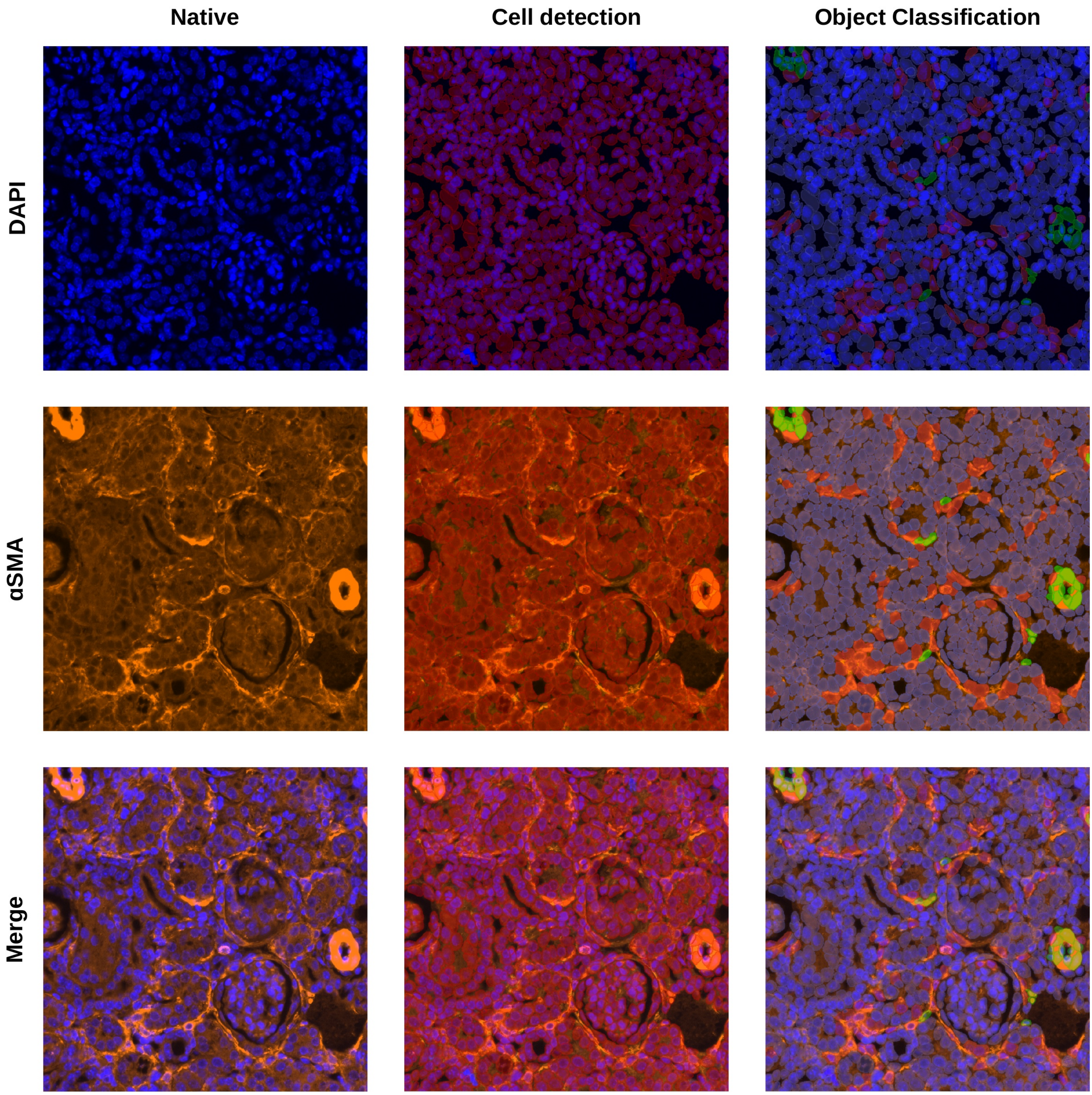


**Supplementary Figure 12**. Example of αSMA-positive interstitial cell detection. Representative images from an obstructed kidney slide immunostained with DAPI and anti-αSMA antibody, illustrating the object classification workflow. Rows show, from top to bottom: DAPI, αSMA, and merged channels. Columns show, from left to right: native image, cell detection overlay, and object classification output. Blue denotes αSMA-negative cells; green denotes vascular smooth muscle cells excluded from quantification (vessel walls) ; red denotes αSMA-positive interstitial cells retained for quantification.
